# Lyophilized mRNA-lipid nanoparticle vaccines with long-term stability and high antigenicity against SARS-CoV-2

**DOI:** 10.1101/2022.02.10.479867

**Authors:** Liangxia Ai, Yafei Li, Li Zhou, Wenrong Yao, Hao Zhang, Zhaoyu Hu, Jinyu Han, Junmiao Wu, Weijie Wang, Pan Xu, Ruiyue Wang, Zhangyi Li, Zhouwang Li, Chengliang Wei, Haobo Chen, Jianqun Liang, Ming Guo, Zhixiang Huang, Xin Wang, Zhen Zhang, Wenjie Xiang, Lianqiang Xu, Bin Lv, Peiqi Peng, Shangfeng Zhang, Xuhao Ji, Huiyi Luo, Nanping Chen, Jianping Chen, Ke Lan, Yong Hu

## Abstract

Advanced mRNA vaccines play vital roles against SARS-CoV-2. However, due to their poor stability, most current mRNA delivery platforms need to be stored at -20°C or -70°C, which severely limits their distribution. Herein, we present lyophilized SARS-CoV-2 mRNA-lipid nanoparticle vaccines, which can be stored at room temperature with long-term thermostability. In the in *vivo* Delta virus challenge experiment, lyophilized Delta variant mRNA vaccine successfully protected mice from infection and cleared the virus. Lyophilized omicron mRNA vaccine enabled to elicit both potent humoral and cellular immunity. In booster immunization experiments in mice and old monkeys, lyophilized omicron mRNA vaccine could effectively increase the titers of neutralizing antibodies against wild-type coronavirus and omicron variants. In humans, lyophilized omicron mRNA vaccine as a booster shot could also engender excellent immunity and had less severe adverse events. This lyophilization platform overcomes the instability of mRNA vaccines without affecting their bioactivity, and significantly improved their accessibility, particularly in remote regions.

## 1. Introduction

After years of research and development, breakthroughs have been made in mRNA delivery systems and mRNA vaccines have become frontrunners that prevent coronavirus disease 2019 (COVID-19). Compared with inactivated vaccines and recombinant protein vaccines, mRNA vaccines could be easily and quickly updated to target different variants, and with comparable first-in-class protective efficacy as well [1]. The sequence-independent manufacturing process saved substantial time and cost in the development of a new vaccine, especially in the pandemic situation, with emergence of new variants of severe acute respiratory syndrome coronavirus 2 (SARS-CoV-2) [2, 3].

Current mRNA therapeutics depend heavily on lipid or lipid-like delivery systems to improve their *in vivo* transfection efficacy [4]. Several lipid components form the nanoparticles carrying the mRNA to targeted organelles or tissues. Although mRNA therapeutics show such superiority, challenges in physiochemical stability still impede their accessibility. Cryogenic preservation and transportation are needed for two current licensed mRNA vaccines, BNT162b2 (−80 ∼-60°C) and mRNA-1273 (−20 °C) [5]. The stringent requirements come from the complex interactions among multiple lipid components and the instability of mRNA, which is sensitive to oxygen, moisture, enzymes, pH and even more [6]. As the exact mechanisms have not been elucidated, it remains a challenge how to improve the stability of mRNA therapeutics [7].

Lyophilization is a process that removes water by sublimation under vacuum at a low temperature [8]. It is a relatively mild drying method and could improve the stability of vulnerable macrobiomolecules or colloidal nanoparticles [9]. Lyophilized mRNA could be stored at 4°C or at room temperature for long time [10, 11]. However, drying of mRNA-lipid nanoparticles (mRNA-LNP) is more sophisticated, as the freezing and dehydration process would introduce mechanical force and deform the vehicle structure, leading to vehicle aggregation, mRNA breakage or leakage [12]. Moreover, some research showed that even though the mRNA-LNP retained their integrity and encapsulation efficiency (EE), the *in vivo* transfection efficacy was greatly reduced after lyophilization due to some unknown causes [13].

Here, we present an optimized lyophilization technique, which could effectively sustain the physiochemical properties and bioactivity of mRNA-LNPs and achieve long-term storage at 2∼8°C and room temperature. The improved thermostability was verified with mRNA-LNPs containing different mRNA molecules, demonstrating its wide applicability. Furthermore, we utilized this technique to prepare the first lyophilized, thermostable mRNA-LNP vaccines encoding the antigen of wild-type (WA1, LyomRNA-WT), delta (LyomRNA-Delta) or omicron COVID-19 variant (LyomRNA-Omicron) and confirmed their high-leveled antibody responses and prevention ability.

Noticeably, we optimized the mRNA sequences in theses lyophilized mRNA vaccines. The lyophilized SARS-CoV-2 vaccines exhibited excellent immunogenicity in mice, rabbits, and old monkeys, and elicited high titers of neutralizing antibodies and potent cellular immune response. In humans, LyomRNA-Omicron given as a booster shot showed excellent performance and could effectively increase the titers of neutralizing antibodies against wildtype coronavirus and many omicron variants.

## 2. Materials and methods

### 2.1 Ethics statement

Mice experiments were conducted by certified staff at the Center for Animal Experiments of Wuhan University and approved by the Institutional Animal Care and Use Committee (AUP #WP2021-0607). Female BALB/c mice aged 6-8 weeks and heterozygous B6/JGpt-H11^em1Cin(K18-ACE2)^/Gpt mice (K18-hACE2 KI mice) aged 5-7 weeks weighing 18-20 g were used. The mice were housed with a 12 h dark-light cycle at a constant temperature (22 ± 2 °C, 45% to 65% relative humidity) and under pathogen-free conditions.

New Zealand rabbits weighing 2∼2.5 kg (half male and female) were purchased from Beijing Longan Experimental Animal Breeding Center. Relative experiments were conducted at Abzymo Biosciences Co., Ltd, which was accredited by Beijing Municipal Science and Technology Commission (SCXK (BJ) 2019-0006).

Nonhuman primate study was performed at Hunan Key Laboratory of Pharmacodynamics and Safety Evaluation of New Drugs.

The protocols and procedures of infectious SARS-CoV-2 viruses under Animal Biosafety Level-III Laboratory facility were approved by the Institutional Biosafety Committee (IBC, Protocol #S01322010B). All samples were inactivated according to IBC-approved standard procedures for removal of specimens from high containment.

For investigator initiated human trials, the protocol and informed consent were approved by Wuhan University and Life Medical Ethics Committee (IRB2022003). Written informed consent was assigned by participants before immunization. This study was conducted in accordance with the Declaration of Helsinki and Good Clinical Practice.

### 2.2 Materials

Lipids were purchased from Xiamen Sinopeg Biotech Co., Ltd. Cholesterol was purchased from AVT (Shanghai) Pharmaceutical Tech Co., Ltd. Ethanol was obtained from Aladdin. Citric acid and trisodium citrate were obtained from Sigma-aldrich. SARS-CoV-2 Wild-type, Delta and Omicron pseudovirus using recombinant replication-deficient vesicular stomatitis virus (VSV) vector that encodes luciferase instead of VSV-G (VSVΔG-Luc) were obtained from Gobond Testing Technology (Beijing) Co., Ltd excepted for these in Extended Data Fig. 12 which were purchased from Vazyme Biotech Co.,Ltd(Nanjing). Omicron-NTD peptide pool (15mers with 11 aa overlap) were synthesized from GL Biochem Co.,Ltd(Shanghai).

### 2.3 RNA synthesis

SARS-CoV-2 NTD-RBD region was used as antigen in this work. The respective N1 -methylpseudouridine modified mRNAs were produced *in vitro* by standard T7 RNA polymerase-mediated transcription reaction and added with Cap1, and then purified though Fast Protein Liquid Chromatography. Luciferase mRNA was synthesized using the same method and purified by MEGAclear™ Transcription Clean-Up Kit (Invitrogen).

### 2.4 Lipid nanoparticle (LNP) preparation and characterization

The mRNA-loaded lipid nanoparticles (mRNA-LNPs) were prepared by mixing an aqueous phase containing mRNA with an ethanol phase containing lipid mixtures using a T-junction mixing device as reported previously [14]. In brief, mRNA was dissolved in citrate buffer (100 mM, pH 4.0). The lipid mixtures were dissolved in anhydrous ethanol at a molar ratio of 46.3:9.4:42.7:1.6 for ionizable lipid, 1, 2-distearoyl-sn-glycero-3-phosphocholine (DSPC), cholesterol, and PEG-lipid. The N/P ratio was kept as 6:1. Then, the ethanol and aqueous phases were mixed at a volume ratio of 3:1 in the T-junction device. Thereafter, mRNA-LNPs were dialyzed against a buffer at pH 7.4 for 6 hours. Then, mRNA-LNPs were sterilized via a 0.22 μm filter and stored at 4°C for further use. The average diameter, polydispersity (PDI) and zeta potential were measured with NS-90Z (Malvern Panalytical). The concentration of leaked mRNA (C_leak_) was determined with a fluorescence detection kit following the manufacture’s protocols. In addition, mRNA-LNP was lysed with 0.4% Triton X-100 to determine the concentration of total mRNA (C_total_). The EE was calculated using the following equation:

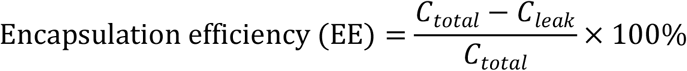

### 2.5 Lyophilization

The mRNA-LNP solution was added with cryoprotectant and filled in a penicillin bottle, and then the mixture was lyophilized with a freezer dryer (Pilot-2H, Boyikang). The resulting powder was collected, characterized, and stored at 4°C for further use. To measure the stability, lyophilized mRNA-LNPs were incubated at either 4, 25, or 40°C (40% humidity) for different time. LNP integrity was measured with a gel retardation assay as previously reported [15]. The mRNA integrity was tested with microfluidic capillary electrophoresis (Agilent Fragment Analyser) [16] and relative mRNA integrity was calculated as % relative to the mRNA-LNP before freeze drying (94% integrity). Lipid composition was analyzed with HPLC-CAD (Thermo Vanquish), with C18 column (Acclaim™, 300Å, 2.1×150 mm), and water (0.5% TEAA) and methanol (0.5%TEAA) as mobile phases.

### 2.6 Cryo-TEM

The cryo-TEM images were obtained from Southern University of Science and Technology. In brief, grids (Quantifoil R 1.2/1.3 300 mesh) were first glow discharged for 60 seconds at 15 mA with a Pelco easiGlow glow discharge unit and 4 μL LNP suspension was applied to the surface of the grid. Grids were then blotted with filter paper (Whatman, Grade 1) for 5.5 s, blot force 0, and after 10 s plunge-frozen in liquid ethane using a Vitrobot Mark IV. The sample chamber of the Vitrobot had a relative humidity of 95% and a temperature of 4 °C, respectively. The grids were imaged using a 300 kV Titan Krios electron microscope (Thermo Fisher Scientific) equipped with a GIF-Quantum energy filter (Gatan) and a slit width of 20 eV was used. Images were recorded with a Gatan K2 direct electron detector operating in super-resolution counting mode at a pixel size of 2.14 Å, with a dose rate of 15 electrons per pixel per second and a total exposure time of 15 s.

### 2.7 Cell culture

Vero E6 (ATCC), HEK 293T/17 (ATCC) and ACE2-expressing 293T cells (Zhejiang Meisen Cell Technology Co., Ltd) were cultured in Dulbecco’s Modified Eagle’s Medium (DMEM) supplemented with 2 mM L-glutamine, 10% FBS (Sigma-Aldrich) and 1% penicillin/streptomycin (BI) at 37°C with 5% CO_2_.

### 2.8 Luciferase transfection *in vivo*

Luciferase mRNA loaded LNPs (mRNA-Luc LNPs) and its lyophilized product, LyomRNA-Luc LNPs were examined with their *in vivo* transfection efficiency. The mRNA-Luc LNP solution (0.5 mL) containing 0.1 mg/mL encapsulated mRNA was freeze-dried with a penicillin bottle. Lyophilized LNPs were reconstituted using 0.5 mL nuclease-free water. Mice were treated with 100 μL mRNA-LNP solution through intravenous (IV) injection into the tail vain. Post 24-hour injection, mice were narcotized with avertin solution (0.2%, 350 μL). Then, the luciferin solution (1.4%, 200 μL) was administered intraperitoneally. The mice were imaged with an IVIS Spectrum *in vivo* imaging system (FluoView400, Boluteng).

### 2.9 Immunization

Three kinds of mRNA-LNPs with mRNA encoding the antigen of wild-type (LyomRNA-WT), delta (LyomRNA-Delta) or omicron (LyomRNA-Omicron) SARS-CoV-2 variant were prepared and used for mouse immunization experiments. The mRNA-LNP solution (0.5 mL) containing 0.1 mg/mL encapsulated mRNA were freeze-dried in penicillin bottles. Lyophilized mRNA-LNPs were reconstituted using 0.5 mL nuclease-free water.

Mice received different doses of freshly prepared or reconstituted LyomRNA-LNPs through intramuscular injections. The immunization procedures and blood collection time intervals varied in different experiments, as indicated in the Results and Discussion.

New Zealand rabbits were immunized twice with different doses (10, 25, 50, 100 μg) of LyomRNA-Omicron through intramuscular injection by a 21 days apart procedure. 14 days after the last shot, neutralizing antibodies titers were detected after blood collection.

5-year-old male rhesus macaques were immunized at Day 0 and Day 21 with 50 μg LyomRNA-Omicron for the primary vaccination. Phlebotomy was conducted at Day 0, Day 7, Day 14, Day 21, and Day 28 followed by serum neutralizing antibodies detection. For the booster scheme, 18 months after two doses of SARS-CoV-2 inactivated vaccine immunization, 50 μg LyomRNA-Omicron were adopted as a booster; Serum neutralizing antibodies were analyzed before and 7, 14, 21, 28 days after boosting.

The blood samples above were collected and centrifuged at 1500 g and 4 °C for 10 min. The supernatant sera were separated, aliquoted, and frozen at -80°C.

### 2.10 Enzyme-linked Immunosorbent Assay (ELISA)

Ninety-six-well ELISA microplates (Greiner) were coated with 2 ng/µL RBD protein (Novoprotein) in coating buffer (Dakewe) at 4°C for 15 h. After washing and blocking, serially diluted mouse sera were incubated in plate at 4°C for 2 h. Following washes, the secondary antibody, goat anti-mouse IgG H&L conjugated HRP (Abcam), was incubated in plate at room temperature for 1 h. Then, 3′5′-tetramethylbenzidine (TMB) (Dakewe) was used as the substrate to detect antibody responses. Data was collected using a microplate reader (Molecular Devices) and SoftMax Pro software version 7.1.0. Endpoint titers were calculated as the dilution that absorbance exceeding 2X background (protein and secondary antibody).

### 2.11 Pseudotyped virus neutralization assay

Neutralizing antibody titers were tested as reported [17]. Briefly, serum samples were diluted and mixed with 1.3×10^4^ TCID_50_/mL of pseudotyped virus and incubated at 37 °C and 5% CO_2_ for 30 min. Thereafter, 293T-ACE2 cells were added and incubated for 24 h. Then, the Bio-Lite luciferase detection reagent (Vazyme) was mixed and incubated for 2 min. The luminescence values (RLU) were detected with the microplate reader immediately, and the amount of pseudotyped virus entering cells was calculated by detecting the expression of luciferase, thus, to obtain the neutralizing antibody content of the sample. Luminescence readout data were collected, and the half maximal inhibitory concentration (IC50) were calculated for the tested samples.

### 2.12 Authentic virus neutralization assay

SARS-CoV-2 Delta-neutralizing antibody was determined by *in vitro* inhibition of cytopathic effect (CPE) and was performed in the Animal Biosafety Level 3 Laboratory at the Center of Laboratory Animal Sciences, Wuhan University (Wuhan, China). Sera from K18-hACE2 KI mice were serially 3-fold diluted in cell culture medium from 1:100 to 1:218700. The diluted samples were mixed with a SARS-CoV-2 Delta virus suspension of 100 plaque-forming units (PFUs), followed by 1 h incubation. The virus-serum mixtures were added to Vero-E6 cells seeded in 24-well plate and cultured in a 5% CO_2_ incubator at 37 °C for 3 days. The inhibitory capacity of each sample dilution was assessed for CPE, and the neutralization titer was calculated as the reciprocal of serum dilution required for 50% neutralization of viral infection.

### 2.13 Intracellular Cytokine Staining

For ICS assays, mice were sacrificed 3 weeks following the second immunization and spleens were harvested. Spleen single cell suspensions were prepared in lymphocyte isolation solution (Dakewe) by mashing tissue against the surface of a 70 µm cell strainer (Falcon). Erythrocytes were removed by density gradient centrifugation. Cells from each mouse were resuspended in R10 medium (RPMI 1640 supplemented with 10% HI-FBS, 1% Pen-Strep solution, 2 mM L-Alanyl-Glutamine, and HEPES) and detected cell viability by Countess 3 Automated Cell Counter (Invitrogen). For mouse intracellular cytokine staining in T cells, 4 × 10^6^ spleen cells were incubated for 7 hours at 37°C in a 5% carbon dioxide incubator with protein transport inhibitor cocktail (4A Biotech) under two conditions: no peptide stimulation (DMSO) and stimulation with Omicron-BA.1 NTD-RBD peptide pools. Peptide pools were used at a final concentration of 4 µg/mL each peptide. Following stimulation, cells were washed twice with PBS buffer and stained with a kit of Zombie Aqua™ Fixable Viability (Biolegend, cat.423101) for 20 min at RT. Cells then washed in cell staining buffer (BD, cat.554657) and resuspended in Fc Block (Biolegend, cat.101320) for 5 min on ice prior to immunostaining with a surface stain cocktail containing the following antibodies: I-A/I-E Alexa Fluor 488(Biolegend, cat.107616), CD8a PerCP/Cyanine5.5 (Biolegend, cat.100734), CD44 Alexa Fluor 700 (Biolegend, cat.103026), CD62L BV650 (Biolegend, cat.104453), CD4 BV605 (Biolegend, cat.100548), and CD3ε APC/Fire 750 (Biolegend, cat.152308) in cell staining buffer (BD). After 30-min incubation, cells were washed twice and incubated with 500 µL BD Fixation and Permeabilization Solution (BD, cat.554722,1X) at 4°C for 20 min, washed twice with BD perm wash buffer (BD, cat.554723), and stained with a cocktail of fluorescently labeled anti-cytokine antibodies: IFN-γ PE/Dazzle 594 (Biolegend, cat.505845), TNF-α PE/Cyanine7 (Biolegend, cat.506324), IL-2 BV421 (Biolegend, cat.503826), IL-4 PE (Biolegend, cat.504104), and IL-5 APC (Biolegend, cat.504306), and CD3ε APC/Fire 750 (Biolegend, cat.152308) in cell staining buffer (BD). After 30 min, cells were washed once with 1X perm wash buffer (BD) and resuspended in 200 µL cell staining buffer (BD) prior to flow cytometric analysis. The fluorescent signals were analyzed by a CytoFLEX LX flow cytometer (Beckman) and analyzed by FlowJo software (Tree Star, Inc). Background cytokine responses to the no peptide condition were subtracted from that measured in the peptide pool for each individual mouse. Statistical comparison between groups was done by using GraphPad Prism 9 software.

### 2.14 ELISpot assays

IFN-γ ELISpot assays of mice were performed with Mouse IFN-γ Precoated ELISpot Kit according to the manufacturer’s instructions (Dakewe). A total of 1 × 10^5^ splenocytes per well was ex vivo restimulated with the Omicron-BA.1 NTD-RBD peptide mix (2 μg/ml per peptide), negative control (DMSO) or positive control (PMA 50 ng/mL+ Innomycin 1μg/mL), respectively.

IFN-γ ELISpot assays of 5 × 10^5^ Human PBMCs per well were performed with Human IFN-γ Precoated ELISpot Kit purchased from Dakewe. Cells were ex vivo restimulated with the Omicron-BA.1 NTD-RBD peptide mix (4 μg/ml per peptide), negative control (DMSO) or positive control (PHA 2.5 µg/mL), respectively. 24 hours after restimulation, Biotinylated antibody and Streptavidin-HRP were added successively after cell lysis.Then AEC Peroxidase Substrate were added and spots counted using an ELISpot plate reader (Mabtech IRIS™ Fluorospot Reader, Mabtech). Spot numbers were evaluated using Mabtech Apex™ software v.1.1.52.121.

### 2.15 SARS-CoV-2 Delta and Omicron challenge in K18-hACE2 KI transgenic mice

SARS-CoV-2 Delta or Omicron challenge was performed in K18-hACE2 KI transgenic mice at the Animal Biosafety Level 3 Laboratory, Wuhan University.

For Delta variant challenge, thirty-six 6-week-old female K18-hACE2 KI transgenic mice were divided into the negative control group (n=6), the blank lipid nanoparticle (blank LNP) control group (n=6), the low-dose vaccine group (n=12, 5 μg/dose) and the high-dose vaccine group (n=12, 10 μg/dose). The latter three groups were intramuscularly immunized with 0.1 mL blank LNPs, 5 μg LyomRNA-Delta and 10 μg LyomRNA-Delta, respectively, at Day 0 (D0) and D14. At D28 (14 days after the second immunization), mice immunized with LyomRNA-Delta or only blank LNPs were intranasally challenged with 2.5×10^3^ PFUs of SARS-CoV-2 Delta viruses. The challenged mice were observed for clinical symptoms and death and weighed on the day of infection and daily thereafter for up to 14 days post infection (dpi). Mice from the blank LNP control group at 7 dpi, six mice from each vaccination group at 7dpi and six mice from each vaccination group at 14dpi were sacrificed under anesthesia. The lung and brain tissues were collected to determine virus titers by qPCR and pathology score by hematoxylin and eosin h&e staining.

For Omicron challenge, thirty-six 6-week-old male K18-hACE2 KI transgenic mice were divided into the control group (n=7), the low-dose vaccine group (n=8, 5 μg/dose) and the high-dose vaccine group (n=8, 10 μg/dose). Contol group were injected with water.Low-dose and high-dose groups were intramuscularly immunized with 5 μg LyomRNA-Omicron and 10 μg LyomRNA-Omicron, respectively, at Day 0 (D0), D21 and D61. At D73, control and vaccination group were intranasally challenged with 1×10^5^ PFUs of SARS-CoV-2 Omicron-BA.1 viruses. The challenged mice were observed for clinical symptoms and death and weighed on the day of infection and daily thereafter. At 7 dpi, all the mice were sacrificed under anesthesia following virus titers and pathological detection in the lung and brain tissues.

### 2.16 Statistical analysis

Data were collected from at least six independent experiments for *in vivo* experiment. All other experiments were performed at least three times independently unless otherwise specified. Statistical analysis was mainly processed using Prism software version 9.0.0 (GraphPad Software Inc., San Diego, CA) and analyzed by Kruskal-Wallis ANOVA with Dunn’s multiple comparisons test or two-sided Mann-Whitney test.Statistically significant differences were considered when p < 0.05.

For pathology score evaluation, as injury such as bleeding and edema of the entire left lung tissue can be observed by H&E staining under low microscope magnification (10×), the evaluation method is to observe injury distribution from 6 independent visual fields, and score them according to the following criteria (injury area/total area):

0 points: No damage;

1 point: < 25% [Mild injury];

2 points: 25% to 50% [Moderate injury];

3 points: 50% to 75%;

4 points: > 75%.

## 3. Results

### 3.1 Lyophilized mRNA-LNPs exhibited high thermostability and bioactivity

Four kinds of mRNA-LNPs used in this study were prepared through a classic T-junction mixing process and then lyophilized [14]. LNPs containing Luc mRNA (mRNA-Luc LNPs), WT mRNA (mRNA-WT LNPs), Delta mRNA (mRNA-Delta LNPs) or Omicron (mRNA-Omicron LNPs) all showed narrow size distribution and high EE, as listed in Table S1. From the cryo-TEM images (Extended Data Fig. 1), we observed uniform, lamellar and vesicular structures of the mRNA-LNPs.

Lyophilization of LNP was difficult due to the vulnerable nature of mRNA and the sophisticated structures and components in the nanoparticles. Here, we adopted an optimized lyophilization process to eliminate LNP damage during drying. After lyophilization, the dried mRNA-LNPs looked like a white fluffy cake (Extended Data Fig. 2) and were readily and rapidly dissolved in water (<10 s). The reconstituted solution was uniform and translucent, just as the freshly prepared mRNA-LNP solutions. The size, PDI and EE of mRNA-LNPs (Table S1) only changed slightly, indicating that the optimized lyophilization process did not change their basic physical properties. The mRNA integrity was also well maintained (>90%, Extended Data Fig.3), demonstrating that the lyophilization process did not cause damage to mRNA structure.

The *in vivo* transfection efficiency of lyophilized mRNA-LNPs was first evaluated with Luciferase mRNA. As seen in Fig. 1a, mice treated with lyophilized mRNA-Luc LNPs (LyomRNA-Luc LNPs) showed comparable total luminescence intensity to those treated with freshly prepared mRNA-Luc LNPs. Next, we investigated the immunogenicity of mRNA-WT LNPs after lyophilization. Mice were immunized with mRNA-WT LNPs or LyomRNA-WT LNPs at D0 and D7 through intramuscular injection, and at D21, the serum was collected. Neutralizing response was tested at this time point through a pseudotyped virus assay (Fig. 1b). There was no significant difference in the titers against wild-type for the groups treated with 5 μg mRNA-WT LNPs (Geometric mean titers, GMTs, 1991) and 5 μg LyomRNA-WT LNPs (GMTs, 1973). These data demonstrated that the lyophilization process did not affect their bioactivity and immunogenicity.

**Fig. 1.**
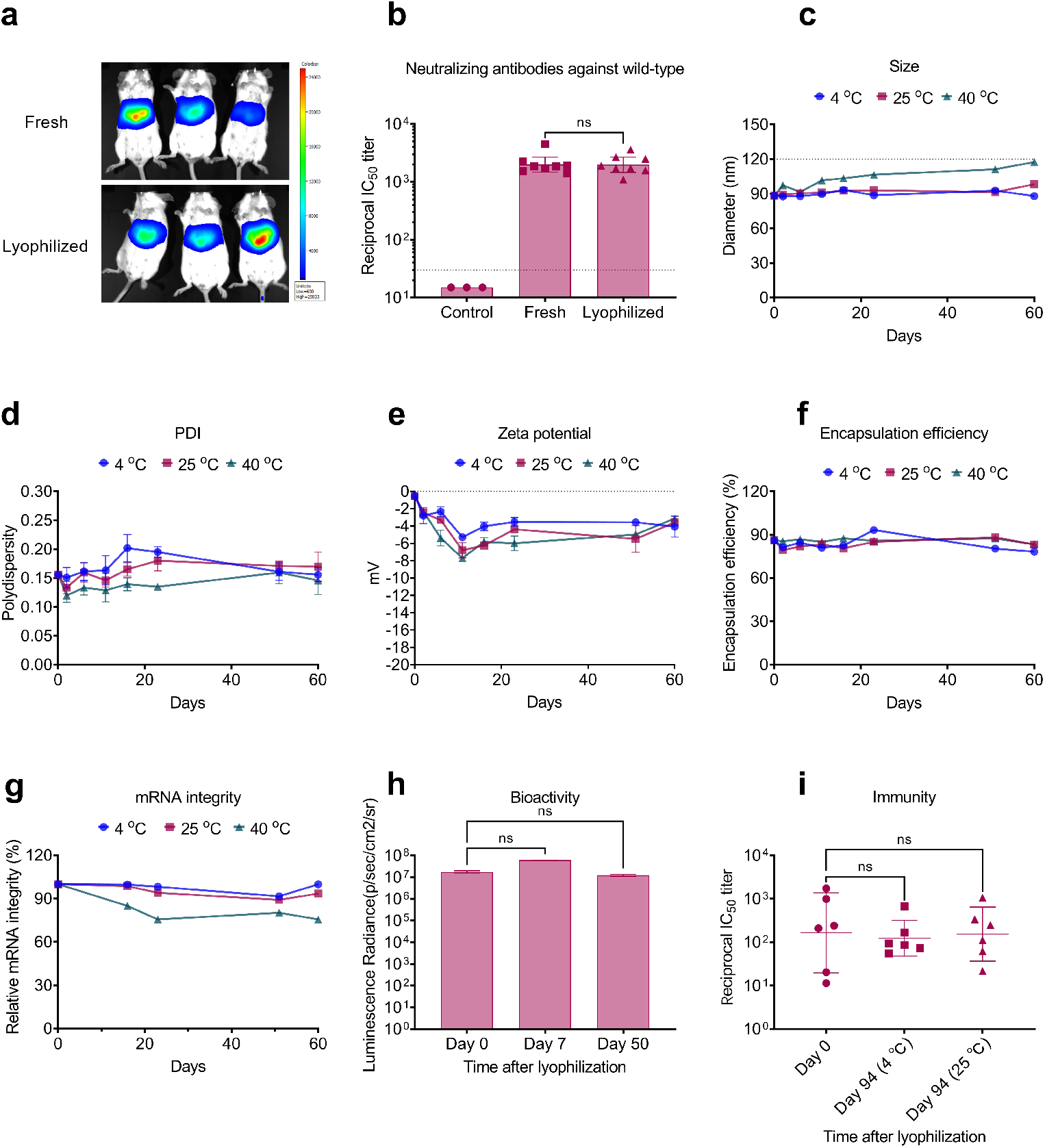
Lyophilized mRNA-LNPs exhibit potent bioactivity and thermostability. (a) Bioluminescent images of mice after treatment with freshly prepared or lyophilized mRNA-Luc LNPs. (b) Pseudotyped IC_50_ titer of mice post two immunizations with freshly prepared mRNA-WT LNPs or LyomRNA-WT LNPs. Mice (n=8) were immunized with mRNA-LNPs (before or after lyophilization) containing 5 μg mRNA at Day 0 and Day 7. The blood was collected and analyzed at Day 21. The dotted line represents assay limit of detection. Change of (c) size, (d) PDI, (e) zeta potential, (f) encapsulation efficiency and (g) mRNA integrity of LyomRNA-Omicron LNPs after incubation at 4, 25, or 40°C over 60 days. (h) Average luminescence radiance of mice after treatment with LyomRNA-Luc LNPs that had been stored for an extended period (n=3). (i) Pseudotyped IC_50_ titer of mice post-two immunizations with freshly prepared mRNA-WT LNPs or LyomRNA-WT LNPs. Mice (n=6) were immunized with mRNA-LNPs (before or after lyophilization) containing 10 μg mRNA at Day 0. Blood was collected and analyzed at Day 14. (b) vaccine groups were compared by two-sided Mann-Whitney test. (h, i) Time points were compared to Day 0 by Kruskal-Wallis ANOVA with Dunn’s multiple comparisons test. ns = not significant. Data are presented as geometric mean ± 95% confidence interval (b, i) or mean with SEM (c-h).

Furthermore, we evaluated the thermostability of mRNA-Omicron LNPs (LyomRNA-Omicron LNPs) by incubating them at 4, 25 or 40°C for different periods of time (Fig. 1c-g, Extended Data Fig. 4). The products did not exhibit obvious change in size, PDI, EE nor mRNA integrity at 4 and 25°C within 94 days (Extended Fig. 4). In the high temperature challenge experiment (40°C), the size of lyophilized mRNA-LNPs increased a little and the relative mRNA integrity remained as high as 75.6% after incubation for 60 days. Furthermore, we analyzed the lipids through high performance liquid chromatograph with Charged Aerosol Detector (HPLC-CAD) after 3-month incubation at 25 °C. As the result showed (Extended Data Fig.5), there was no sign of lipid degradation. The well-maintained physical properties of lyophilized mRNA-LNPs come from the ultra-low water content and oxygen content in the sealing bottles.

The bioactivity was then re-evaluated after long-term storage. The LyomRNA-Luc LNPs after storage at 4°C for 50 days showed no difference from freshly prepared Luc LNPs in the *in vivo* bioluminescence experiment (Fig. 1h). More importantly, after incubation for 94 days at 4 or 25°C, LyoOmicron-LNPs still showed high immunogenicity, which induced pseudotyped IC_50_ titers (Fig. 1i) and IgG titers (Extended Data Fig. 4f) comparable to freshly prepared mRNA-Omicron LNPs.

These results indicated that mRNA LNPs maintained their bioactivity well after lyophilization and the lyophilized product exhibited much improved stability. It could be estimated that the lyophilized mRNA-LNPs could be stored at a 2∼8°C or room temperature for an extended period of time.

### 3.2 High immune response induced by lyophilized mRNA-Delta vaccine

Since 2021, more SARS-CoV-2 variants have continued to emerge, of which Delta and Omicron are the variants with the greatest impact. The fast mutation rate and highly contagious nature of these variants caused heavy casualties and decreased the prevention ability of existing vaccines [18, 19]. Under this circumstance, mRNA vaccines with rapid upgrading ability showed their advantages. For better prevention of the epidemic disease, two mRNA vaccine were prepared to further evaluate their efficacy against Delta and Omicron variants based on the lyophilized mRNA-LNP formulation (LyomRNA-Delta and LyomRNA-Omciron).

Mice received two immunizations with LyomRNA-Delta at an interval of 14 days. Fourteen days after the second immunization, serum samples were obtained to examine the titers of the binding antibodies and neutralizing antibodies. The results showed that the geometric mean of the titers of binding antibodies of the 5 μg and 10 μg group reached 7,980,000 and 5,040,000, respectively (Fig. 2a). Meanwhile, the geometric mean of the titers of neutralizing antibodies against true viruses reached 5978 and 9323, respectively (Fig. 2b), indicating that LyomRNA-Delta elicited very potent humoral immune response.

**Fig. 2.**
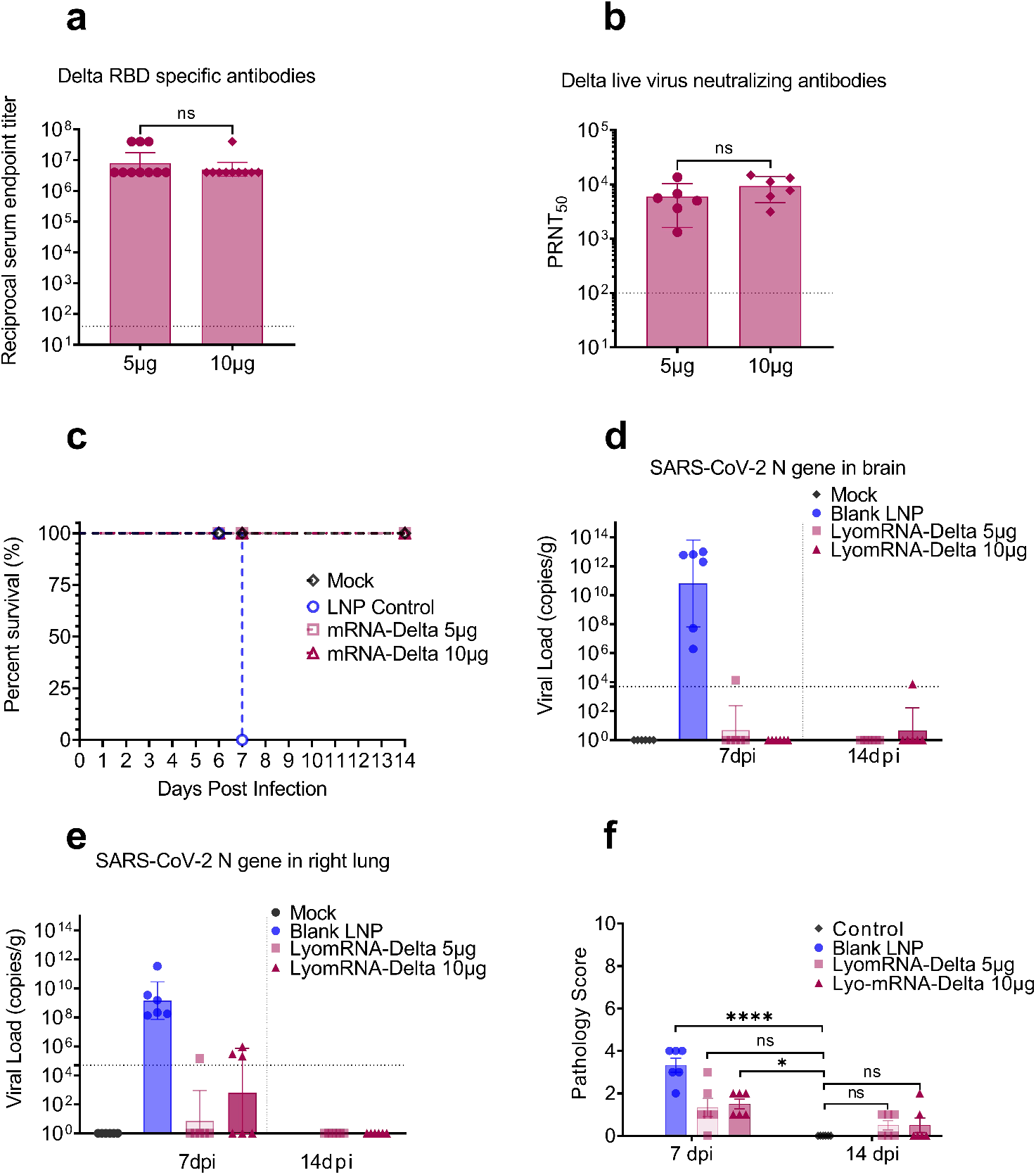
LyomRNA-Detla could elicit potent immune response and protect mice from infection by Delta viruses. (a) BALB/c mice (n=10) were intramuscularly immunized twice at a dose of 5 μg or 10 μg at an interval of 14 days. Plasma samples were collected at Day 14 post immunization for detection of Pseudovirus-based VSV-SARS-CoV-2 Delta 50% neutralization titers. (b-f) K18-hACE2 KI transgenic mice were intramuscularly inoculated with 5 μg or 10 μg LyomRNA delta vaccine twice, separated by 14 days, and two weeks after the second dose, 2.5×10^3^ PFUs of SARS-CoV-2 Delta were administered to the animals intranasally. The mock group was immunized with saline. The blank-LNP group was immunized with lipid nanoparticles without loaded mRNA. (b) Live VSV-SARS-CoV-2 Delta 50% neutralization titers 14 days after the second dose. (c) Animal survival curve after challenge, all blank LNP animals died or were euthanized 7 days after challenge. Virus load in the brain (d), virus load in the lungs (e) and pathology score (f) were evaluated 7 and 14 days after challenge. (a-b) Doses were compared by two-sided Mann-Whitney test. (f) Groups were compared to mock by Kruskal-Wallis ANOVA with Dunn’s multiple comparisons test. ns=not significant, * = p-value < 0.05, *** = p-value < 0.001. Data are presented as geometric mean ± 95% confidence interval (a-b), geometric mean ± geometric SD (d-e) or mean with SEM (f). The dotted line represents assay limit of detection.

After receiving two doses of 5 μg and 10 μg LyomRNA-Delta vaccine, K18-hACE2 KI mice were challenged with SARS-CoV-2 Delta infection. The group received blank LNPs were highly susceptible to SARS-CoV-2 Delta infection. Beginning at 2 dpi, mice demonstrated clinical symptoms, including lethargy, ruffled fur, arched back, and drowsiness. At 6 dpi (days post infection), body weight reduction was seen in the blank LNP control mice, and approximately 20% weight loss was found by 7 dpi, and two mice died, and the remaining mice were moribund and humanely euthanized according to ethic principle. In contrast, mice that received two doses of reconstituted LyomRNA-Delta showed no clinical abnormality, with no difference in body weight from the mock mice (Fig. 2c, Extended Data Fig. 6a). For these groups, six mice were euthanized at 7 dpi, and the remaining six mice were euthanized at 14 dpi. After euthanasia, brain and right lung lobes were collected from each mouse for viral load quantification, and the left lung lobes were fixed for histopathological evaluation. The results showed that lyophilized mRNA-Delta vaccination successfully protected the mice from SARS-CoV-2 Delta infection. The viral load in the right lung tissue of LyomRNA-Delta vaccinated mice remained at the level of the low limit of quantification at 7 dpi. In contrast, a high level of viral load was detected in the right lung tissue of mice in the blank LNP control group, with up to approximately 10^11^ copies/g (Fig. 2d). Importantly, the viral load in the right lung tissue was almost undetectable in all LyomRNA-Delta vaccinated mice at 14 dpi (below the limit of detection). The viral load in the brain tissues in the 5 μg groups displayed similar patterns as those in the lung tissues. Additionally, LyomRNA-Delta 10 μg showed remarkable protective effect on virus dissemination and amplification (Fig. 2e). Moreover, pathological study showed at 7 dpi, the vaccinated mice showed markedly less severe lung tissue injuries than the blank LNP group, with no difference in the pathology score from the mock mice. At 14 dpi, the lung tissues of the vaccinated mice showed further improvement(Fig. 2f, Extended Data Fig. 6b). Together, these data suggest that lyophilized mRNA-Delta vaccine was highly immunogenic and could fully protect the challenged mice from SARS-CoV-2 Delta infection.

### 3.3 Lyophilized Omicron mRNA vaccine could simultaneously elicit high levels of humoral immunity and cellular immunity

The Omicron variant has become the predominantly prevalent SARS-CoV-2 strain worldwide. Consequently, we specifically developed the freeze-dried mRNA vaccine LyomRNA-Omicron based on the sequence of the Omicron variant. In mice, a scheme consisting of two immunizations at an interval of 21 days led to a positive antibody conversion rate of 100% after the first shot of LyomRNA-Omicron. At D21, the geometric mean titer of neutralizing antibodies reached 1064 in the group of 1μg dose. The titer of neutralizing antibodies continued to climb after the 2^nd^ immunization and was 23-fold higher at D35 than that at D21 and the geometric mean titer reached 24256. Meanwhile, the plasma levels of neutralizing antibodies correlated positively with the dosage; however, there was no statistical difference in the dose range from 1 ug to 5 ug. In addition, the titer of binding antibodies for Omicron RBD showed an identical trend (Fig. 3a). Furthermore, in the binding antibody typing experiment, the LyomRNA-Omicron activated Omicron-specific RBD IgG2a/IgG ratio was greater than 1 (Extended Data Fig. 8a).

**Fig. 3.**
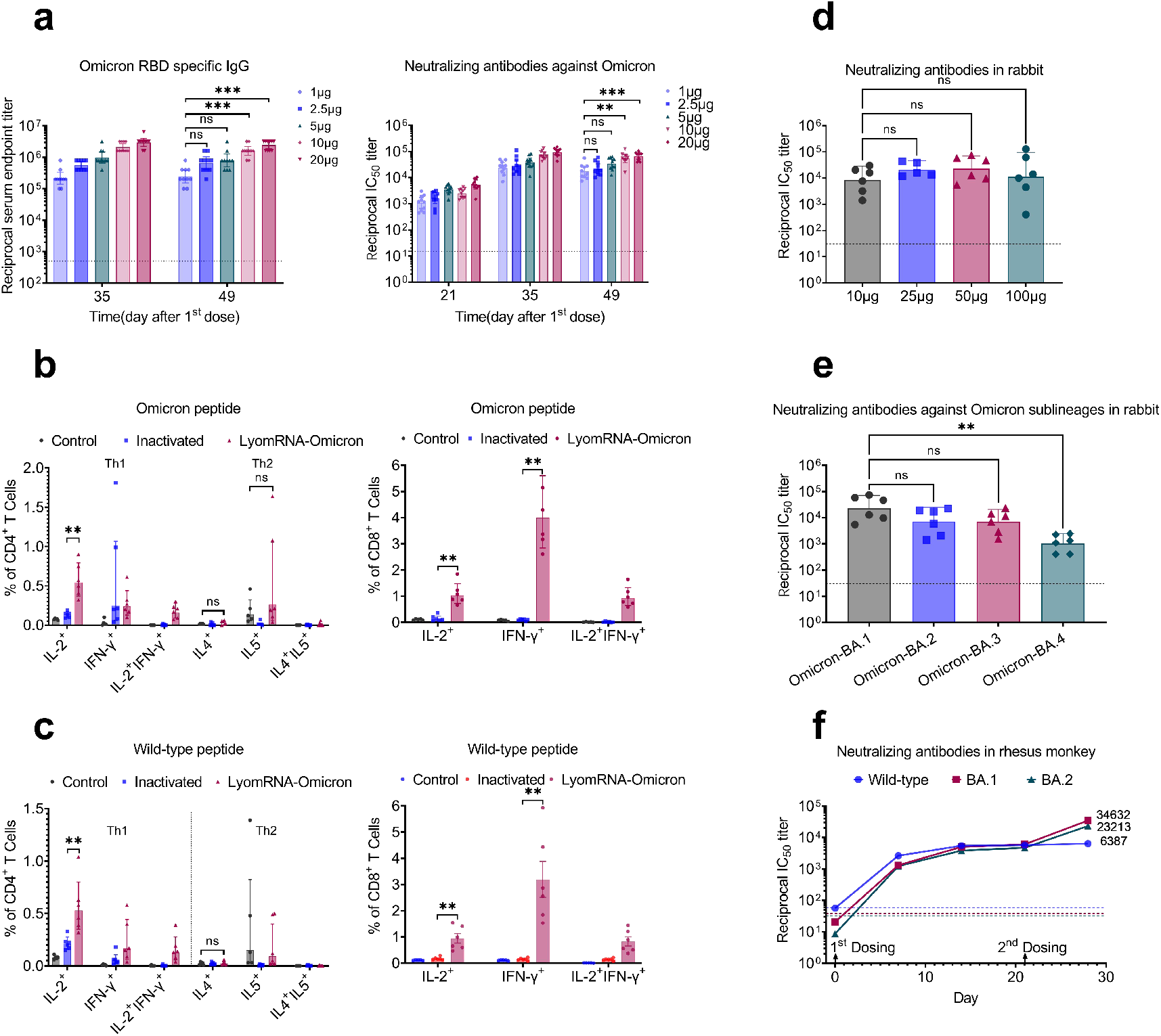
LyomRNA-Omicron could simultaneously activate high levels of humoral and cellular immunity. (a-b) C57BL/6N mice (n=) received two muscular injections of different doses of LyomRNA-Omicron, the time for the 1^st^ dose was considered Day 0, and the 2^nd^ dose was given 21 days after the1^st^ dose. Plasma Omicron RBD specific IgG binding antibody titers at different time points following the 1^st^ dose(a) and the titers of pseudovirus neutralizing antibodies (b). (c-d) C57BL/6N mice were vaccinated with LyomRNA-Omicron 5 μg or inactivated vaccine 0.65 U at an interval of 21 days. The spleens were obtained 28 days after the second shot for ICS. The control mice received normal saline. The frequencies of IL2/IFN-γ/IL4/IL5 positive CD4 T cells and IL2/IFN-γ positiveCD8 killer cells specific for SARS-CoV-2 Omicron NTD-RBD peptide (c) or wild-type NTD-RBD peptide (d) were detected. (e-f) New Zealand rabbits received two muscular injections of vaccines at an interval of 21 days. The plasma neutralizing antibodies for pseudoviruses (e) and pseudoviruses of all Omicron variants at the dose of 50 μg (f) were detected 14 days after the last immunization. (g) Five-year-old monkeys were immunized each with 50 μg LyomRNA-Omicron (n=3), twice at a 3-week interval. Blood samples were collected before or after vaccination. Neutralizing antibodies against the wild-type, Omicron variant BA.1 and BA.2 were tested by a VSV-pseudovirus based system. Immunogenicity data at 7, 14, 21 and 28 days after the first dose of LyomRNA-Omicron were collected. (a-b, e-f) Group comparisons were made by Kruskal-Wallis ANOVA with Dunn’s multiple comparisons test. (c-d) Vaccine groups were compared by two-sided Mann-Whitney test. ns=no significant, ** = p < 0.01, *** = p < 0.001. Data are presented as geometric mean ± 95% confidence interval (a-b, e-g) or geometric mean ± geometric SD (c-d). Dotted lines represent assay limits of detection for (a-b, e-f) and plasma pseudovirus neutralizing antibody titers of control monkey without immunization for (g).

We also examined LyomRNA-Omicron cellular immunity by ICS. The results showed that, in C57 and Balb/c mice stimulated by Omicron NTD-RBD peptide pool, LyomRNA-Omicron could effectively activate IL2^+^, IFN-γ^+^ Th1 CD4 cells and the ratio of activated CD4 cells was higher than that with inactivated vaccine(CoronaVac, inactivated whole wild-type virus vaccine by Sinovac Life Sciences). Meanwhile, there was no statistical difference in activated IL4^+^, IL5^+^ Th2 CD4 cells compared with the PBS control group. In the same time, LyomRNA-Omicron also activated the production of massive amounts of CD8^+^ cytotoxic T cells against Omicron, and the ratio of activated cells was also significantly higher than that of the inactivated vaccine group (Fig. 3b, Fig. 8b, Extended Data Fig. 10a, Extended Data Fig. 11a/b/c). Intriguingly, LyomRNA-Omicron could also effectively activate cellular immune response against SARS-CoV-2 wild-type viruses (Fig. 3c, Extended Data Fig. 10b).

We examined immune response to LyomRNA-Omicron in rabbits. A scheme consisting of two immunizations at an interval of 21 days was used and the levels of neutralizing antibodies were detected at D14. The results showed that the geometric mean titer reached 8337 with 10 μg LyomRNA-Omicron, and there was no statistical difference in antibody levels in the dose range from 10 to 100 μg (Fig. 3d). Meanwhile, LyomRNA-Omicron could effectively neutralize Omicron BA.1/BA.2/BA.3 subvariants. Though it had certain neutralizing activities against BA.4, the geometric mean titer was reduced by 20 folds compared that for BA.1 (Fig. 3e).

The elderly population is most vulnerable to SARS-CoV-2. Therefore, we used old monkeys to evaluate the level of immunity against LyomRNA-Omicron. In the primary immunization experiment with 2 immunizations at an interval of 21 days, 50 μg LyomRNA-Omicron simultaneously elicited the production of neutralizing antibodies against Omicron-BA.1, Omicron-BA.2 and the wild-type viruses in old monkeys (Fig. 3f).

In summary, LyomRNA-Omicron could elicit effective humoral immunity and Th1-dominant cellular immunity.

### 3.4 Lyophilized Omicron mRNA vaccine could offer immune protection as a booster shot in animals

Multiple key mutations in Omicron allow the viruses to escape immune protection of two-dose immunization with vaccines that were designed based on the sequence of the original sequence of the virus [20]. Therefore, booster immunity is critical for warding off Omicron. So far, at least two thirds of the world’s population have received at least one shot of SARS-CoV-2 vaccine, and inactivated vaccine is one of the most commonly administered vaccine especially in China [21]. Therefore, we designed LyomRNA-Omicron based on inactivated vaccine (CoronaVac, inactivated whole wild-type virus vaccine by Sinovac Life Sciences) to carry out heterogeneous booster experiment.

In mice, the third immunization with LyomRNA-Omicron following two immunizations with inactivated vaccine elicited high levels of neutralizing antibodies against the wild-type, Delta, Omicron-BA.1, and Omicron-BA.2 variants simultaneously and the geometric mean titer increased by 9, 13.5, 50.7, and 77.7 folds over that before vaccination, respectively. Meanwhile, the titer was significantly higher than that with the third booster shot with heterogeneous inactivated vaccine and was 3.4, 5.7, 46, and 43.3 times higher than the latter, respectively (Fig. 4a/b/c/d). In 5-year old monkeys, heterogeneous booster with LyomRNA-Omicron 18 months after primary immunizations with two-dose inactivated vaccines could elicit the production of high levels of neutralizing antibodies against the wild-type, Omicron-BA.1 and Omicron-BA.2 variants, with a geometric mean titer of 6803, 2454, and 1046, respectively (Fig. 4e), indicating that though the levels of neutralizing antibodies against SARS-CoV-2 were low in primates long after immunization with inactivated vaccine, booster immunization with LyomRNA-Omicron could still engender high levels of neutralizing antibodies.

**Fig. 4.**
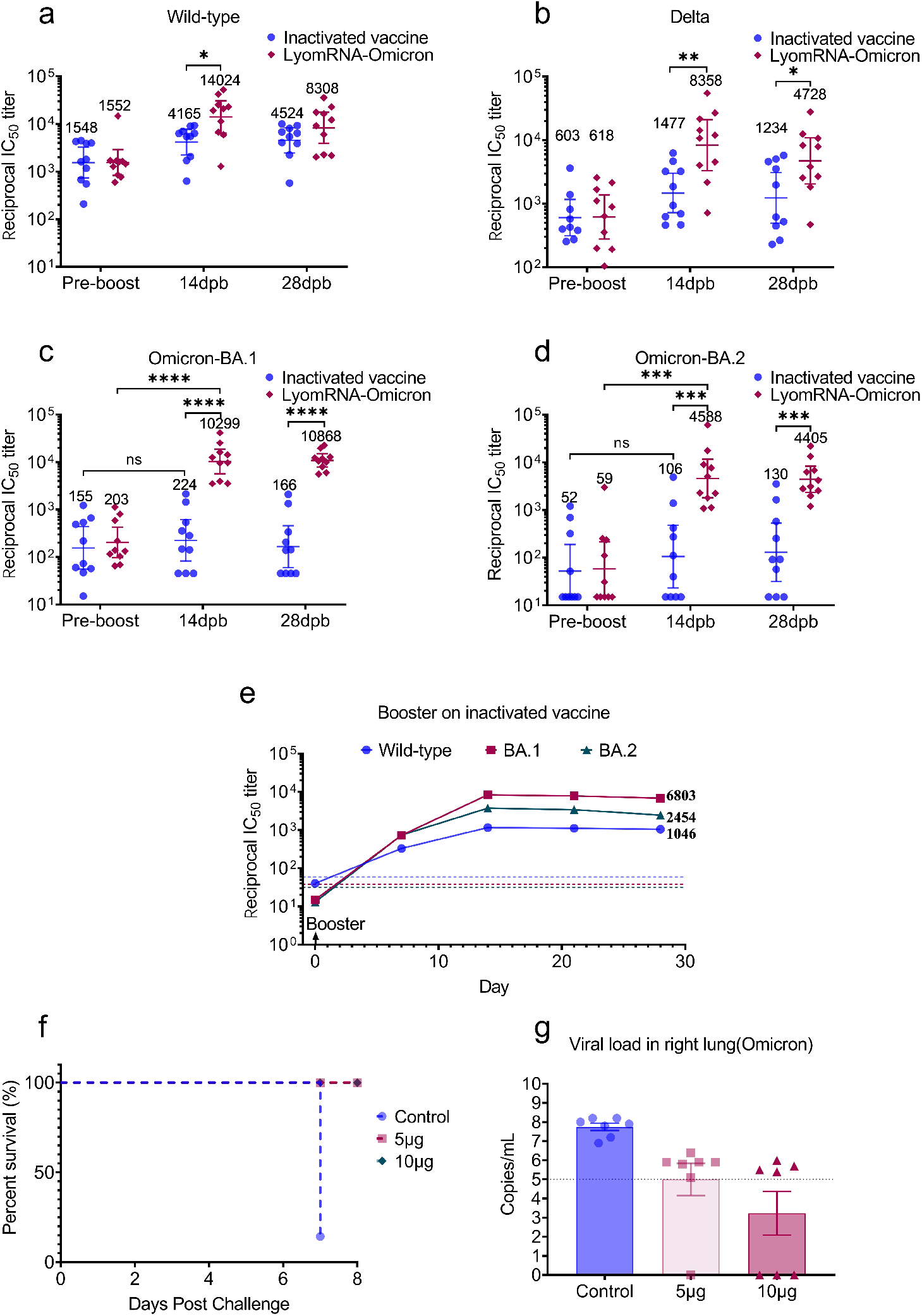
LyomRNA-Omicron booster immunization engenders excellent immune protection. (a-d) The titers of plasma neutralizing antibodies against wild-type (a), Delta (b), Omicron-BA.1 (c), and Omicron-BA.2 (d) pseudoviruses were detected in BALB/c mice (n=10) at day 0 and 21 of immunization with inactivated vaccine (60 SU per dose), day 42 of immunization with heterogeneous LyomRNA-Omicron (5 μg dose) or homogeneous inactivated vaccine (60 SU dose), before booster immunization and 14 (14 dpb) and 28 days post booster immunization (28 dpb). (e) Five-year-old monkeys that had received two doses of inactivated vaccines (50 μg dose per animal, twice at a 3-week interval) were boosted 18 months later with LyomRNA-Omicron 50 μg. Blood samples were collected before or 7, 14, 21, and 28 days after boosting with LyomRNA-Omicron. Neutralizing antibodies against the wild-type viruses, Omicron variant BA.1 and BA.2 were tested by a VSV-pseudovirus based system. Dotted lines represent blood pseudovirus neutralizing antibody titers of non-immunized control monkeys. (f-g) K18-hACE2 KI transgenic mice were vaccinated at day 0, 21, and 61 at a dose of 5 μg and were lethally challenged at day 73 with Omicron-BA.1. Survival (f) and lung virus load at day 7 post challenge (g) were recorded. (a-d) The groups were compared by two-sided Mann-Whitney test. (f) Within each dose level by two-sided Wilcoxon signed-rank test, and doses were compared post-boost by Kruskal-Wallis ANOVA with Dunn’s multiple comparisons test. (d-f) Vaccine groups were compared by two-sided Mann-Whitney test. * = p < 0.05, ** = p < 0.01, *** = p < 0.001, **** = p < 0.0001. (a-e) Data are presented as geometric mean ± 95% confidence interval(a-e) or mean with SEM (g).

In addition, we designed homogeneous booster experiment with LyomRNA-Omicron in *hACE2* transgenic mice. Two primary shots were done at D0 and D21 followed by the third booster shot at D73. The mice were lethally challenged with Omicron-BA.1. The results showed that the survival rate of mice immunized with low dose (5 μg) and (10 μg) reached 100%, and 6/7 non-immunized control mice died at D7 post challenge (Fig. 4f). The viral load of the right lung and pathological section of the left lung revealed that compared to non-immunized mice, the immunized mice were apparently protected against Omicron (Fig.4g, Extended Data Fig. 7).

The data demonstrated that in animal studies, LyomRNA-Omicron as a booster showed remarkable immunization effects.

### 3.5 Lyophilized Omicron mRNA vaccine booster immunization experiment in humans

Most people in China have received inactivated SARS-CoV-2vaccine. Booster immunization is a feasible approach to prevent breakthrough infection by Omicron. We tested the immunization effect of LyomRNA-Omicron as a booster shot in humans.

26 volunteers participated in the experiment; 19 of them had received two shots of inactivated vaccines for more than 6 months since the last immunization. They were assigned to Group I, that is, heterogeneous booster with LyomRNA-Omicron based on previous two-dose immunization with inactivated vaccines. The remaining 7 participants had received 3 inoculations with inactivated vaccines for 2 months since the last shot. They were assigned to Group II, that is, LyomRNA-Omicron as a heterogeneous booster based on previous three-dose immunizations with inactivated vaccines.

In Group I, the pre-booster plasma samples of the subjects contained very low tiers of neutralizing antibodies against the wild-type strain, with a geometric mean titer of 27, and the titer was nearly 0 for Omicron-BA.1 and Omicron-BA.2. Fourteen days after boosting with 50 μg LyomRNA-Omicron, the geometric mean titers of neutralizing antibodies against the wild-type, Omicron-BA.1, and Omicron-BA.2 all significantly climbed, reaching 6827, 5800 and 4196, respectively, and rising by 253, 725 and 420 folds. The high levels of antibodies were maintained at 1- and 2-months post boost. Among them, the geometric mean titer of antibodies at D28 post heterogeneous boosting with LyomRNA-Omicron was significantly higher than that of the control group (using inactivated vaccine as the third shot), and was 18.5, 64.6, and 39.6 folds higher for the wild-type, Omicron-BA.1, and Omicron-BA.2 compared with the latter (Fig. 5a/b/c, Extended Data. Fig.13a).

**Fig. 5.**
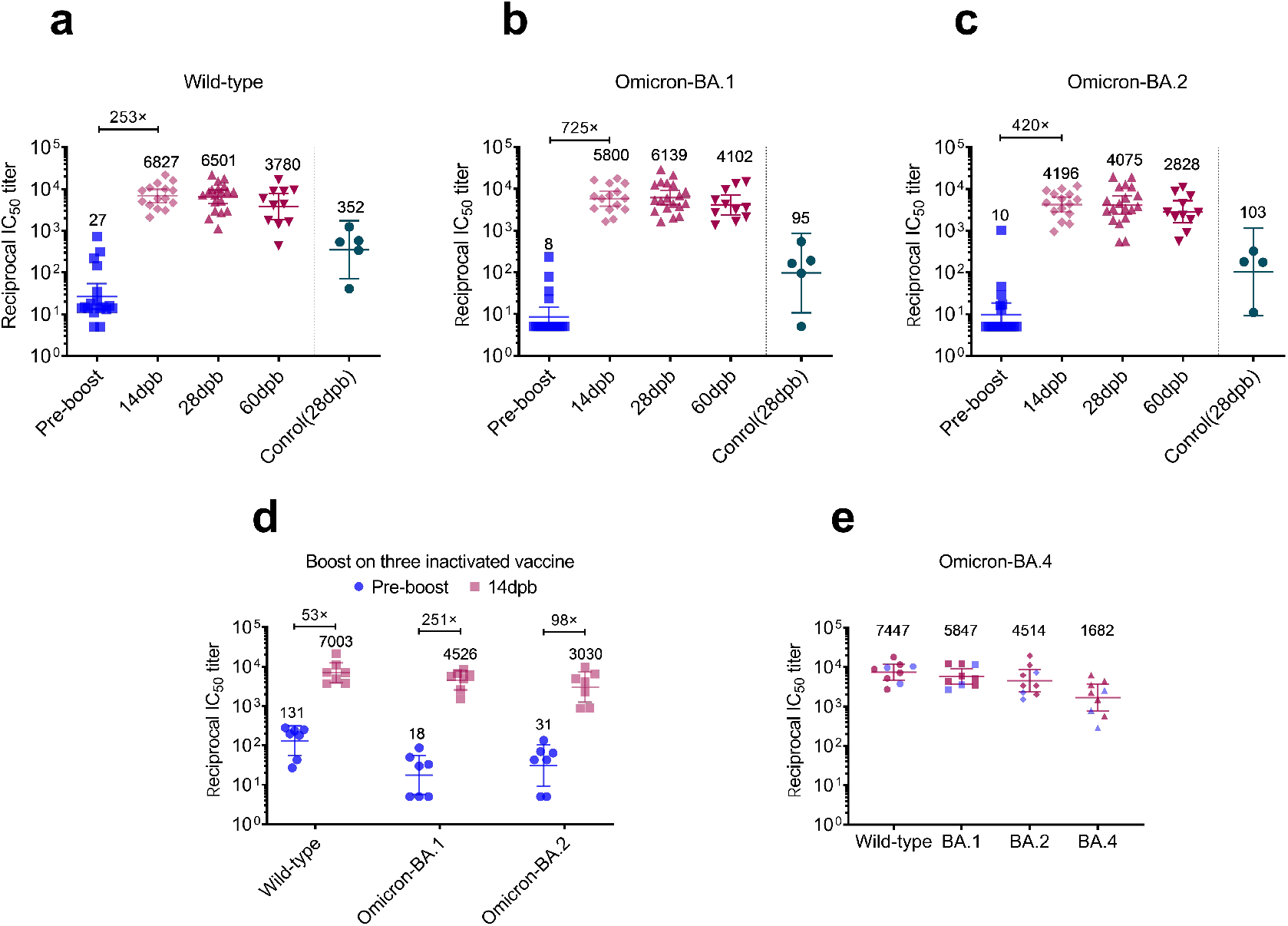
Booster immunization in humans with LyomRNA-Omicron could elicit high titers of neutralizing antibodies. (a-c) On the basis of two doses of inactivated vaccine (at least 6 months from the last dose), a booster shot with LyomRNA-Omicron was given. Then, the tiers of neutralizing antibodies against (a) the wild-type, (b) Omicron-BA.1 and (c) Omicron-BA.2 pseudoviruses were detected before boost (day 0, pre), 14 (14 dpb) and 60 days post booster (60 dpb) were detected. Plasma neutralizing antibodies in the control group one month after booster immunization with homogeneous inactivated vaccine. (d) A 4^th^ booster with LyomRNA-Omicron was given on the basis of 3 immunizations with inactivated vaccine (after at least 6 months post the final dose), and then the titers of plasma neutralizing antibodies against the wild-type, Omicron-BA.1, and Omcron-BA.2 pseudoviruses 14 days post booster immunization. (e) The titers of plasma neutralizing antibodies against Omicron-BA.4 14 days after LyomRNA-Omicron booster immunization (blue represents booster based on 2 inoculations with inactivated vaccine and red represents booster based on 3 inoculations. Data are presented as geometric mean ± 95% confidence interval.

In Group II, LyomRNA-Omicron was used as the fourth booster shot, and the geometric mean titers of neutralizing antibodies at D14 post the booster shot against the wild-type, Omicron-BA.1, and Omicron-BA.2 were 7003, 4526 and 3030, respectively, and were increased by 131, 251 and 98 folds versus those before the booster shot. However, by only looking at the data, the titers of neutralizing antibodies were comparable for heterogeneous booster with LyomRNA-Omicron following two or three inoculations with inactivated vaccines (Fig. 5d, Extended Data. Fig.13b).

Currently, Omicron-BA.4\BA.5 are more infectious and spreading quickly. We found that in humans receiving booster immunization with LyomRNA-Omicron on the basis of inactivated vaccine, the geometric mean titer of neutralizing antibodies against BA.4 was reduced by mild 4.4-fold than wild-type, and 3.5 folds than that for BA.1, with the latter consistent with previous reports [22, 23]. Interestingly, this is different from the 20-fold steep reduction for BA.4 compared to BA.1 in rabbits, reflecting immune difference between humans and animals. However, the levels of neutralizing antibodies against BA.4 were still high after boosting with LyomRNA-Omicron, with the geometric mean titer reaching 1682 (Fig. 5e).

In addition, booster immunization with LyomRNA-Omicron could elicit potent cellular immune response to Omicron variant (Extended Fig. 13).

No grade 2 adverse events were observed in the 26 volunteers who received LyomRNA-Omicron and no one had a body temperature above 37.5°C. 4 persons reported apparent symptoms, including redness and swelling in the muscular injection site in 1 subject, dizziness in 1 subject, and pain in the axillary lymph nodes in 2 subjects, all of which disappeared within 5 days.

In summary, LyomRNA-Omicron is very suitable as a booster shot based on inactivated vaccines and could elicit excellent immune effects on the SARS-CoV-2 wild-type and Omicron variants and at the same time has mild side effects.

## 5. Conclusion

The mRNA-LNP vaccine has several advantages, such as rapid design of antigen, potential for quick, inexpensive, and scalable manufacturing, and inducing strong humoral and cellular immunity, which make it a powerful weapon against infectious diseases, especially against those caused by viruses that are easy to mutation. The advantage and efficacy of mRNA-LNP vaccines has also been proved by two approved and marketed vaccines against COVID-19, Comirnaty and Spikevax.

According to the data in Our World in Data as of July. 11, 2022, 66.8% of the world population has received at least one dose of a COVID-19 vaccine. Totally 12.15 billion doses have been administered globally. However, only 20% of the population in low-income countries have received at least one dose of vaccine [21]. It is mainly because of the instability of mRNA-LNP vaccines, which require ultracold temperature (−80°C∼-60°C) for storage and transportation, greatly limiting their accessibility. Therefore, it is particularly important to develop a mRNA-LNP vaccine that can be transported in conventional cold-chain for broad application.

In this study, we successfully achieved the long-term storage of mRNA-LNP vaccine through lyophilization. The optimized lyophilization process preserved the nanostructures and physiochemical properties of mRNA-LNP and well maintained its bioactivity, showing no significant difference from freshly prepared vaccine. Moreover, lyophilized mRNA-LNPs exhibited a much-improved thermostability at 4 and 25°C, without obvious deterioration after incubation for 52 days.

The lyophilized mRNA-WT, mRNA-Delta and mRNA-Omicron vaccine could both produce high levels of IgG or neutralizing antibodies. In the challenge study of LyomRNA-Delta vaccine against Delta strain, it was confirmed that the vaccine could fully protect mice from infection and clear the virus. In the two-dose primary vaccination scheme, LyomRNA-Omicron elicited immune response by Th1 biased CD4 T cells and CD8 killer T cells against both Omicron and the wild-type strain. Besides, It could also induce a wide range and high-level immune response against Wild-type, Delta and Omicron variants as a booster for inactivated vaccines both in mice and old monkeys, with better efficacy than inactivated vaccine. In human studies, LyomRNA-Omicron as a booster shot on the basis of inactivated vaccine caused fewer side effects, without a single case of fever. Following the booster shot, the production of neutralizing antibodies against Omicron can be elicited 100% and the titers of neutralizing antibodies against the wild-type, Omcron-BA.1 and BA.2 strains were increased by 100 folds versus those before the booster shot. Furthermore, faced with the rapidly spreading Omicron-BA.4, the titers of neutralizing antibodies only decreased 3.5 folds compared to those against BA.1, but remained at an elevated level.

As LyomRNA-Omicron adopts an advanced mRNA-LNP freeze-drying process, making it more thermostable and accessible, it is in the ranks of advanced and excellent vaccine by having fully addressed the storage and transportation issue of current mRNA-LNP vaccines.

In conclusion, the lyophilized mRNA-LNP vaccine has the advantage of both high immunogenicity and accessibility and is very suitable for prevention of epidemic such as SARS-CoV-2.

## Acknowledgment

We thank Dr. Zan Tang for guidance on histopathology analysis. We thank Prof. Zheng Liu, Xiaoyu Ma and Jiamin Gu for conducting cryo-TEM experiments. We thank Dr.Tao Pang for advice on antibodies detection and animal studies.

## Author contributions

Y. Hu, K. Lan, and J. Chen conceptualized and supervised the whole study. Y. Hu and L. Ai conceived, planned, designed the strategy and experiments. L.Ai conducted and supervised RNA synthesis, mRNA-LNP preparation and animal studies. Y. Li developed, conducted, and optimized LNP preparation, lyophilization procedures and mRNA-LNP analysis. Z. Li, H. Zhang and W. Yao planned and supervised animal experiments and human trials. Z. Hu, J. Han, J. Wu, H. Chen, and J. Liang conducted LNP preparation, lyophilization and analysis. R. Wang and Z. Li performed cell experiment and pseudotyped virus neutralization assay. Z. Li measured IgG titers. W. Wang, P. Xu, B. Lv, P. Peng, and S. Zhang conducted and helped modified the mRNA synthesis and LNP preparation procedures. C. Wei, M. Guo, Z. Huang, X. Wang, Z. Zhang, W. Xiang, and H. Luo helped conducted the animal experiments. L. Xu and N. Chen conducted the HPLC-CAD analysis.Y.Hu, L.Ai, Y.Li wrote the manuscript. All authors supported the review and editing of the manuscript.

---

**Table S1.** Main physical characteristics of mRNA-LNPs before and after lyophilization.

| Samples | Fresh |  |  | Lyophilized |  |  |
| --- | --- | --- | --- | --- | --- | --- |
|  | Size (nm) | PDI | EE (%) | Size (nm) | PDI | EE (%) |
| mRNA-Luc LNPs | 74.1 | 0.19 | 94.7 | 93.9 | 0.15 | 85.6 |
| mRNA-WT LNPs | 92.4 | 0.14 | 92.1 | 117.7 | 0.16 | 87.0 |
| mRNA-Delta LNPs | 72.8 | 0.03 | 93.2 | 96.0 | 0.10 | 85.6 |
| mRNA-Omicron LNPs | 77.6 | 0.09 | 94.3 | 88.4 | 0.16 | 86.2 |

**Extended Data Fig. 1.**
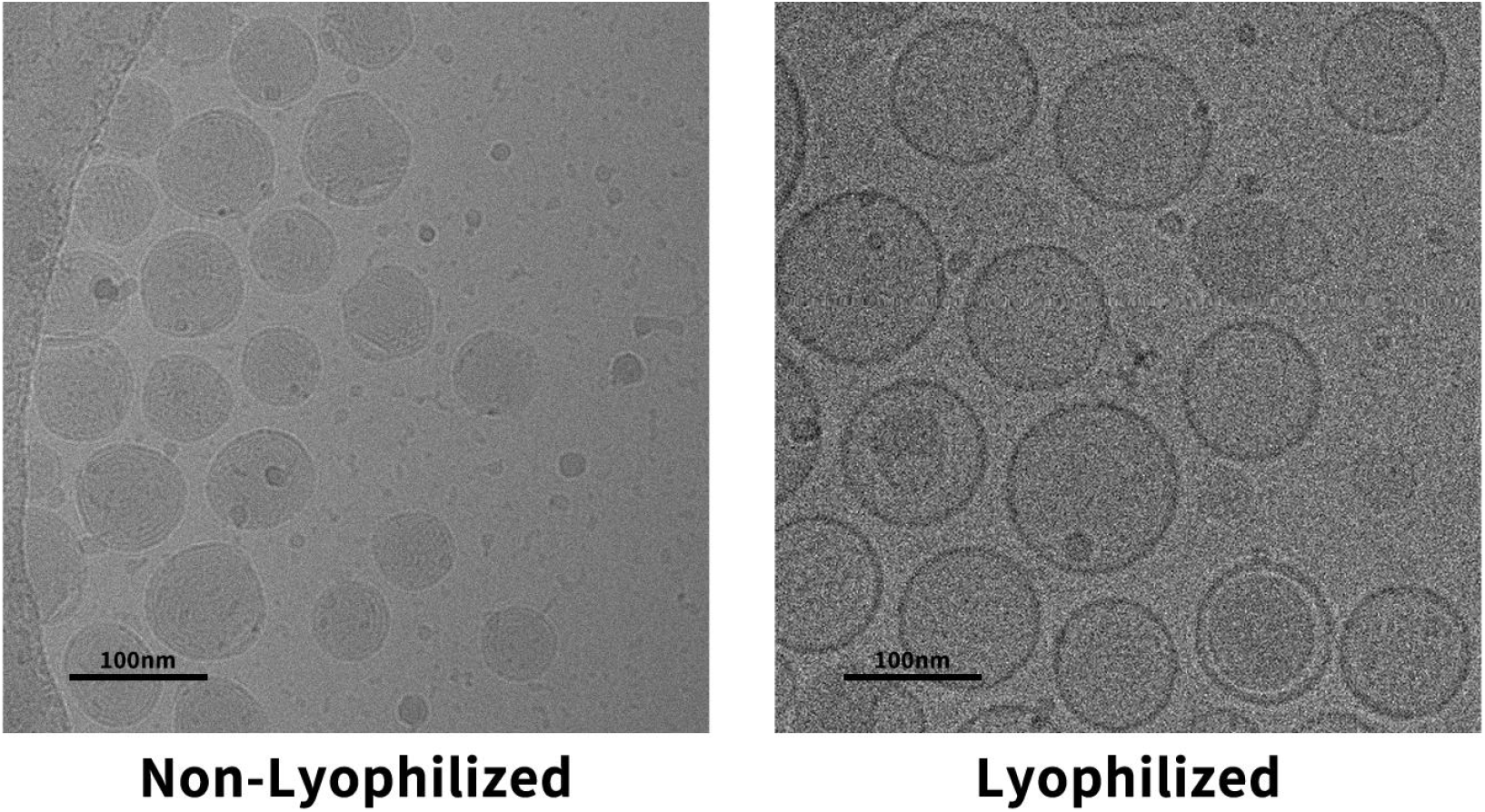
Cryo-TEM image of fresh or lyophilized mRNA-LNPs.

**Extended Data Fig. 2.**
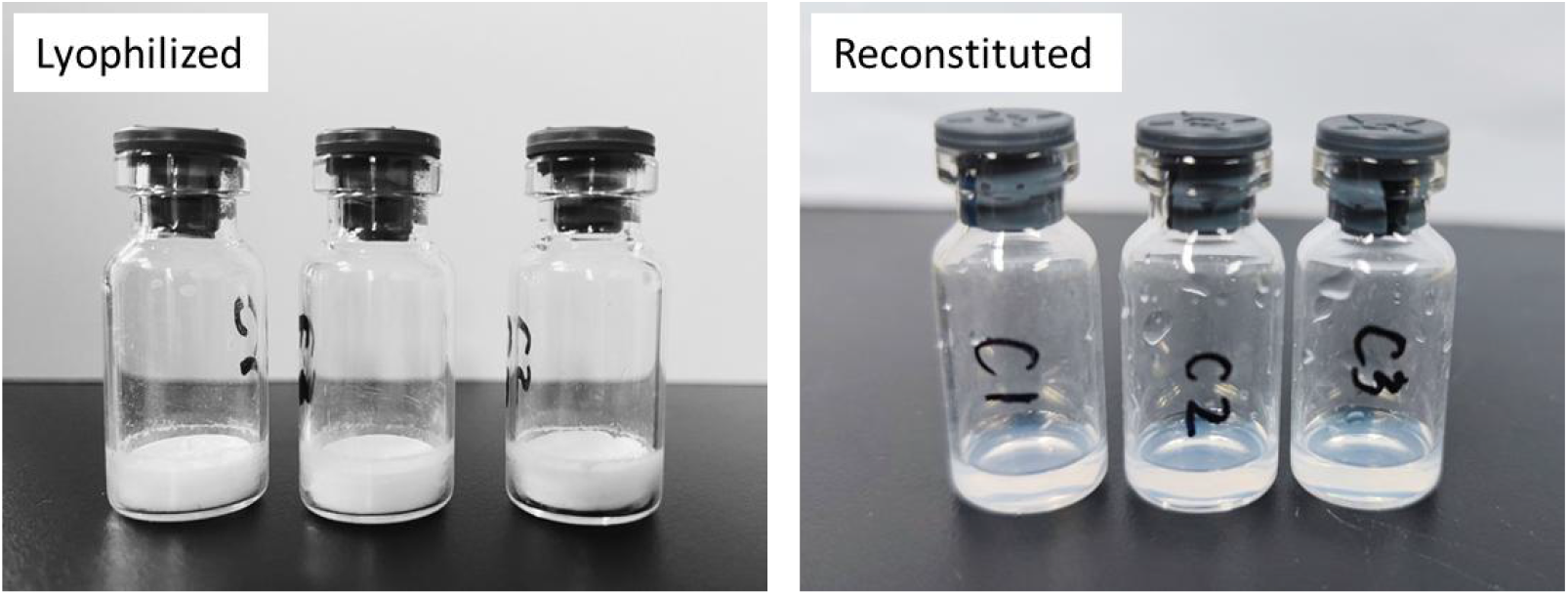
Images of lyophilized mRNA-LNPs and the reconstituted solutions.

**Extended Data Fig. 3.**
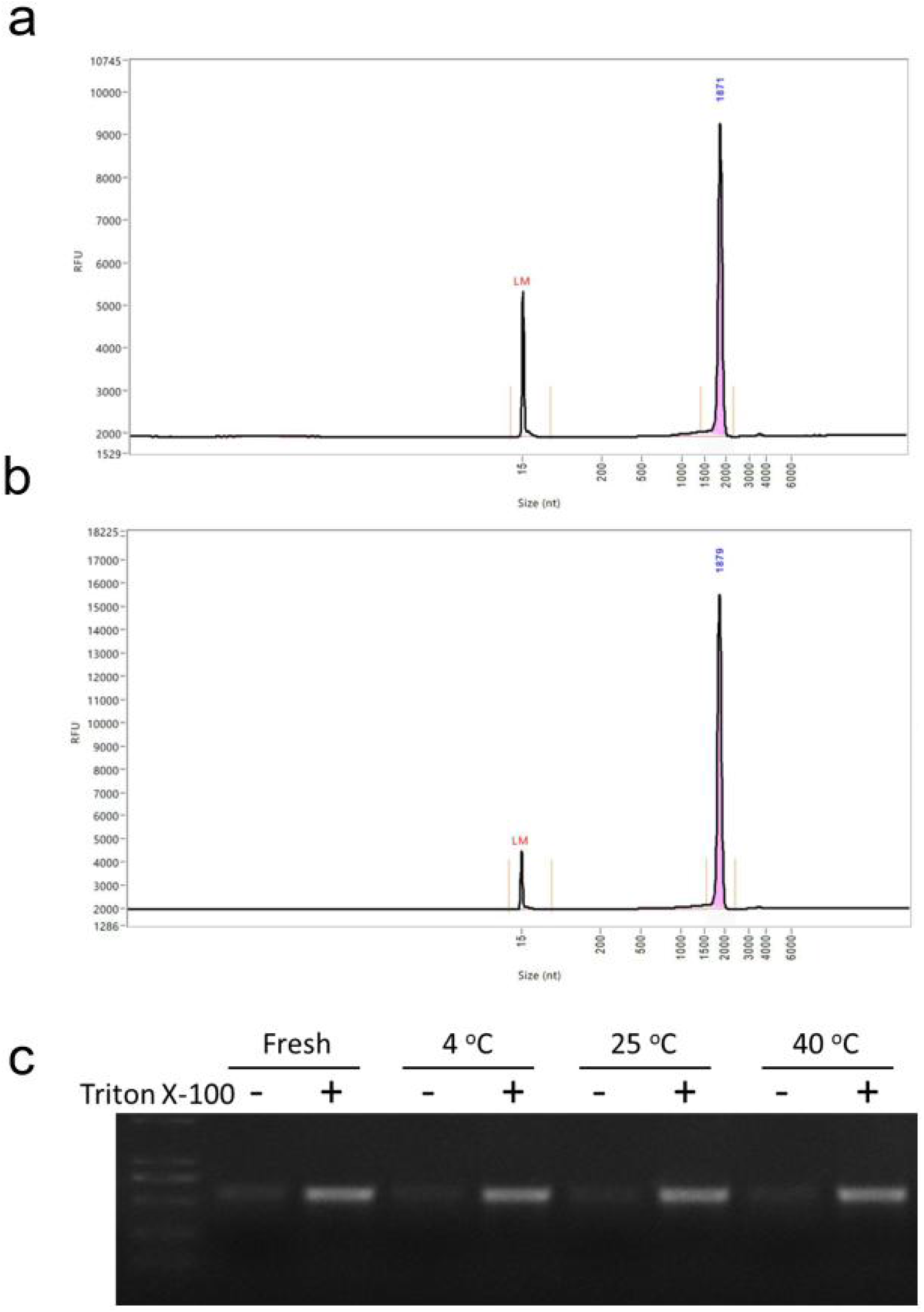
Microfluidic capillary electrophoresis analysis of (a) Omicron mRNA and (b) reconstituted LyomRNA-Omicron. The mRNA integrity was 94.1% and 93.9%, respectively. (c) gel electrophoresis images of LyomRNA-Omicron LNPs after incubation at 4, 25, or 40°C for 10 days.

**Extended Data Fig. 4.**
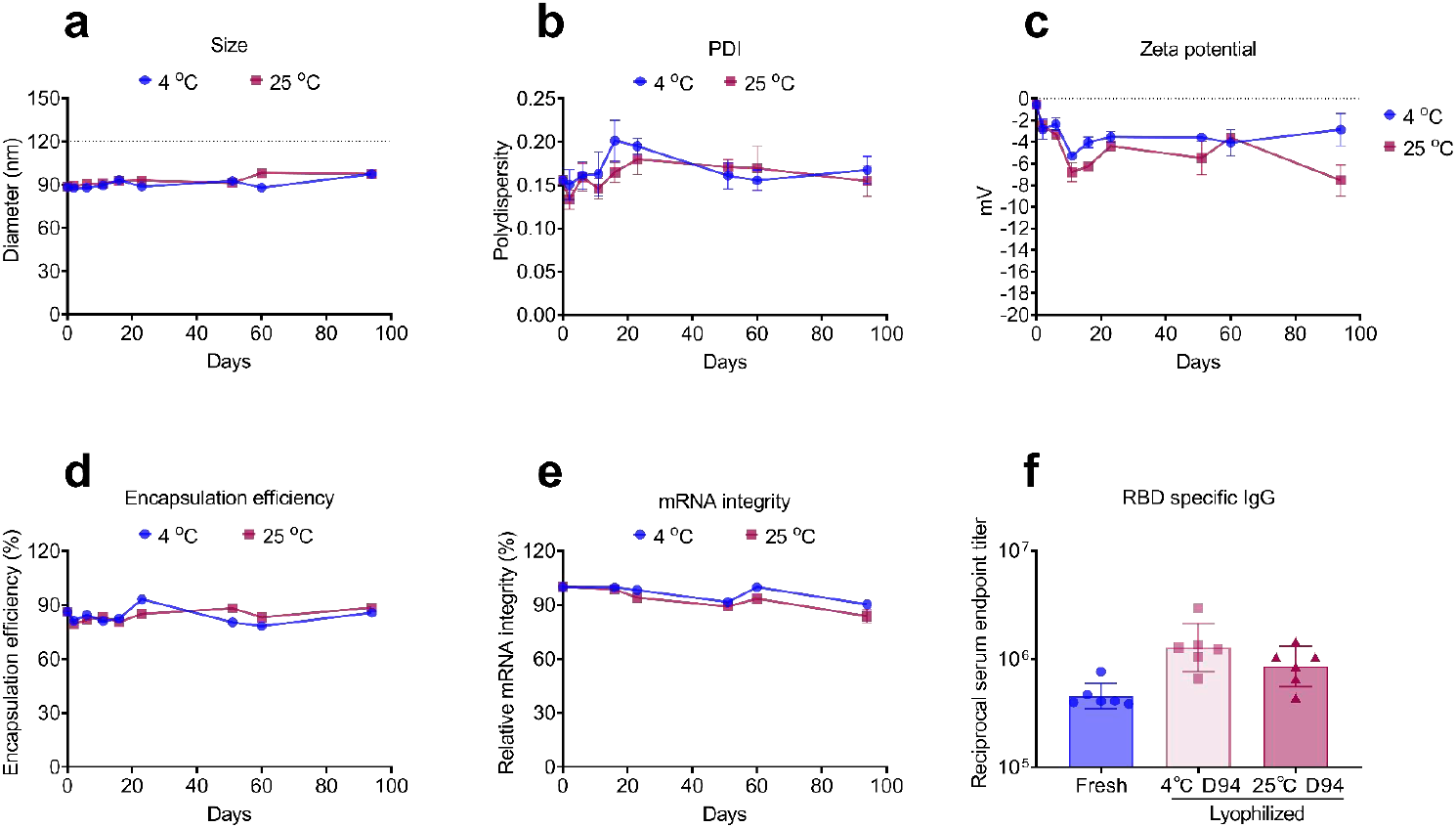
Long term stability of freeze-dried mRNA-LNP. Change of (a) size, (b) PDI, (c) zeta potential, (d) encapsulation efficiency and (e) mRNA integrity of LyomRNA-Omicron LNPs after being incubated at 4, 25 or 40 °C over 94 days. (f) Total IgG titer of mice (n=8) post two immunizations with freshly prepared mRNA-Omicron LNPs or Lyo mRNA-Omicron LNPs. Mice were immunized with mRNA-LNPs (before or after lyophilization) containing 10 μg mRNA at day 0. The blood was collected and analyzed at day 14. Data are presented as geometric mean ± 95% confidence interval(a-e) or mean with SEM (f).

**Extended Data Fig. 5.**
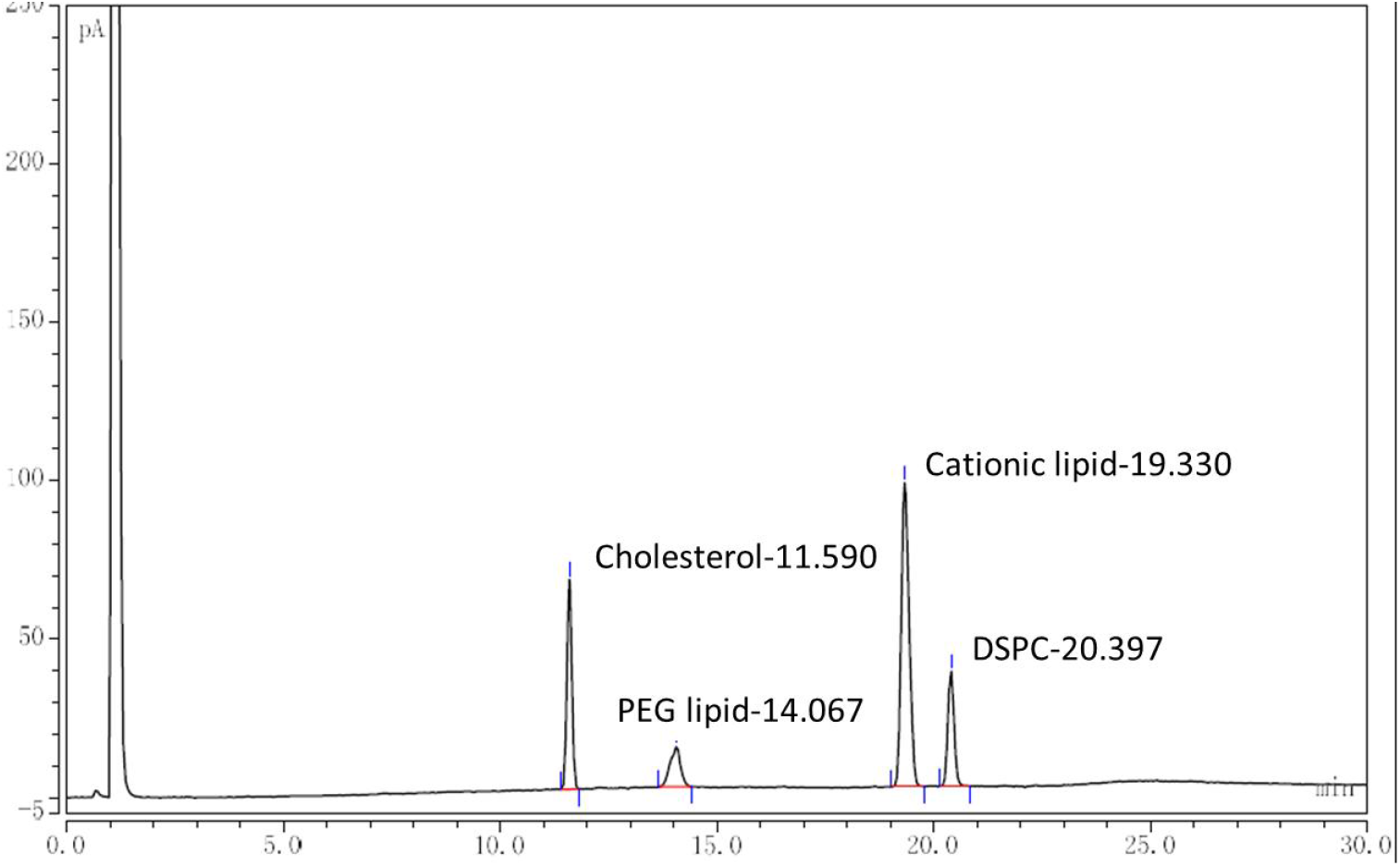
High chemical stability of Lyo-Omicron-mRNA-LNPs. Lyo-Omicron-mRNA-LNPs were incubated at 25 °C for 3 months, then the lipid components were analyzed with HPLC-CAD.

**Extended Data Fig. 6.**
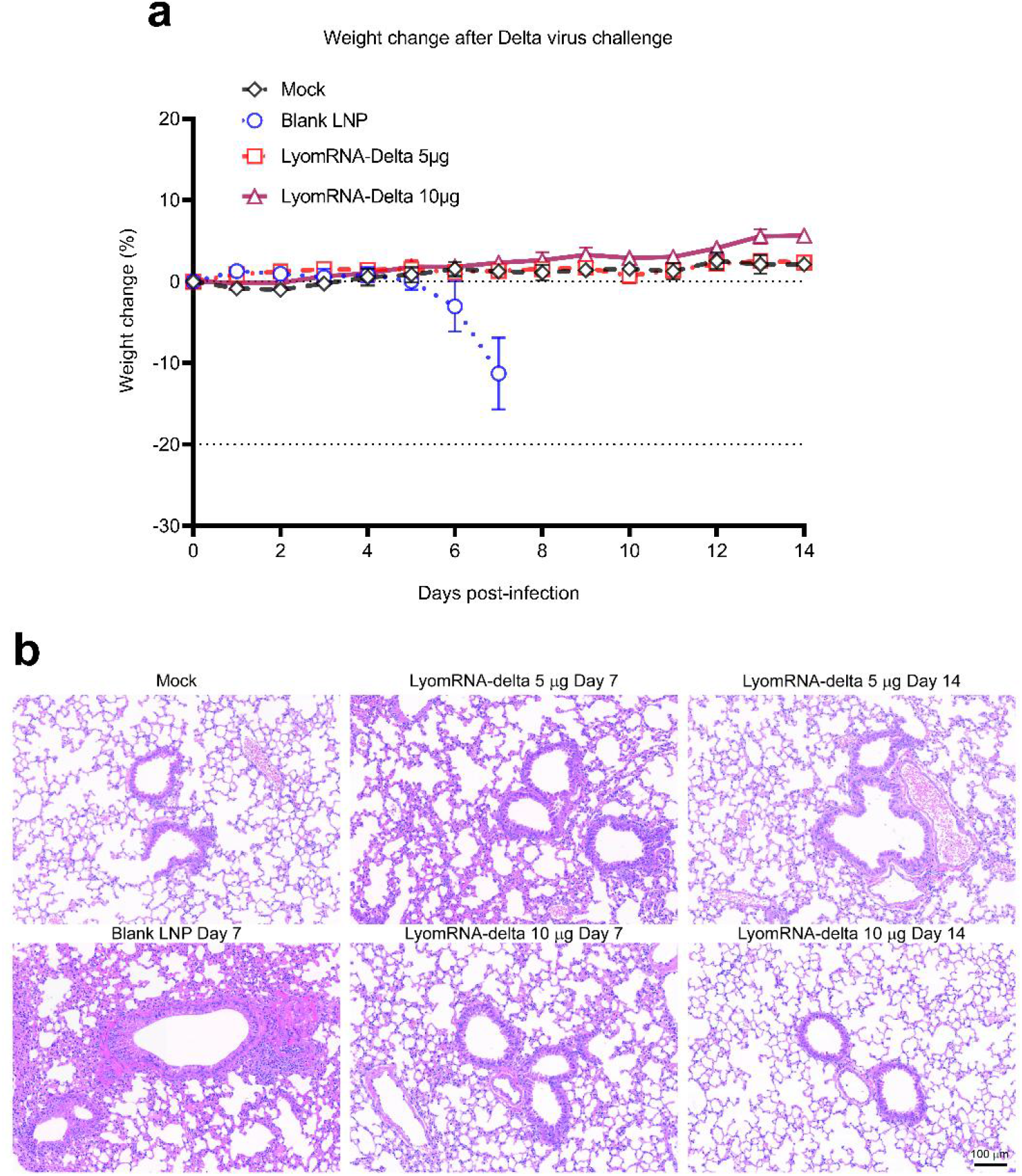
Detla virus challenge experiment. (a) Changes in body weight of mice post virus challenge. (b) Hematoxylin and eosin-stained lung sections were examined at day 7 and day 14 after challenge. Data are presented as mean with SEM.

**Extended Data Fig. 7.**
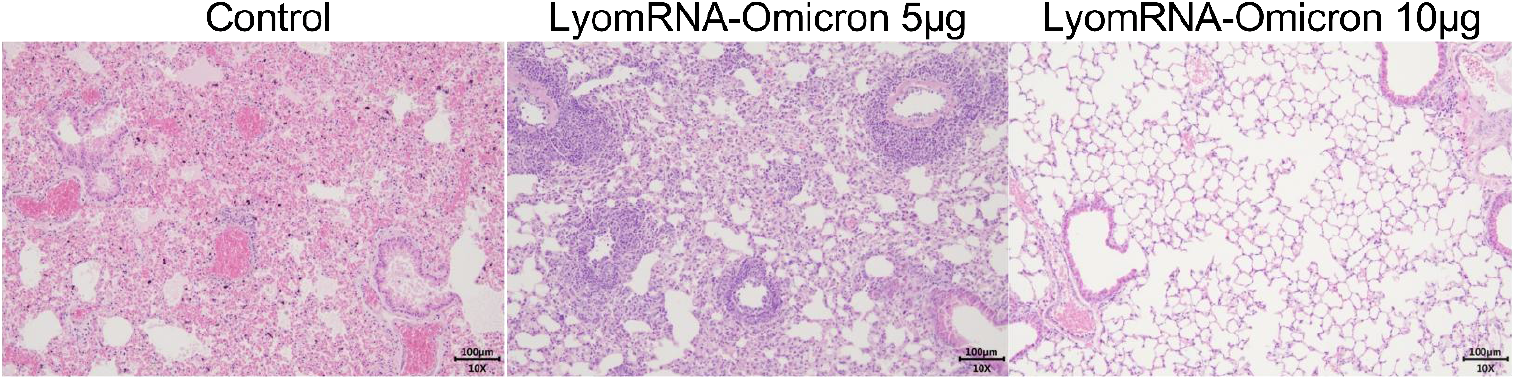
K18-hACE2 KI transgenic mice were immunized at a dose of 5 μg at day 0, 21, and 61 and were lethally challenged at day 73 with Omicron-BA.1. Pathological sections were evaluated 7 days post challenge.

**Extended Data Fig. 8.**
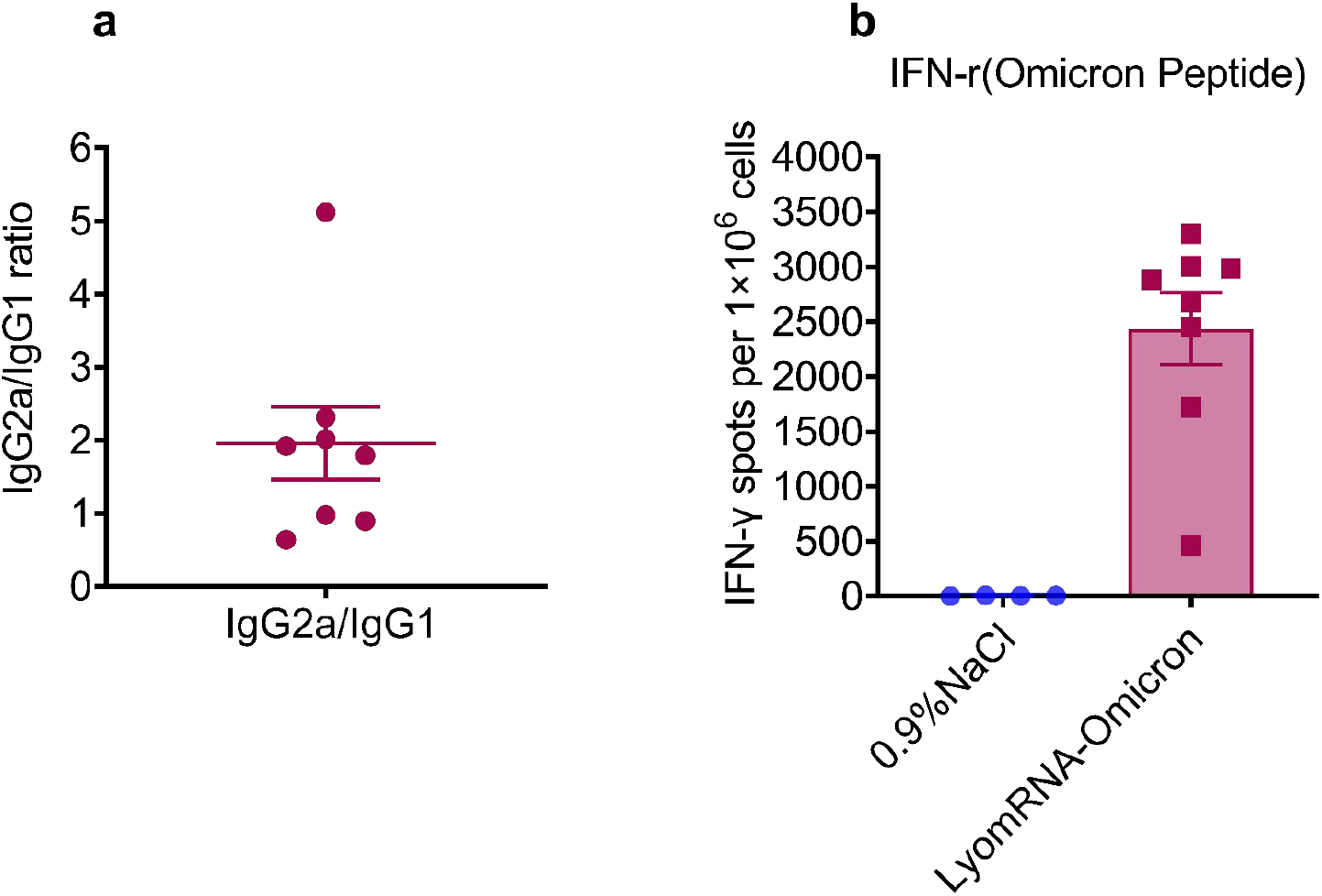
Mouse immunogenicity of LyomRNA-Omicron. (a)BALB/c (n=8) received the second immunization with 5 μg LyomRNA-Omicron at an interval of 7 days. The ratio of plasma Omicron RBD-specific IgG2a/IgG1 was detected at day 14 post immunization. (b) The number of IFN-r+ spots in mouse spleen one month after immunization was detected using ELISpot. Data are presented as mean with SEM.

**Extended Data Fig. 9.**
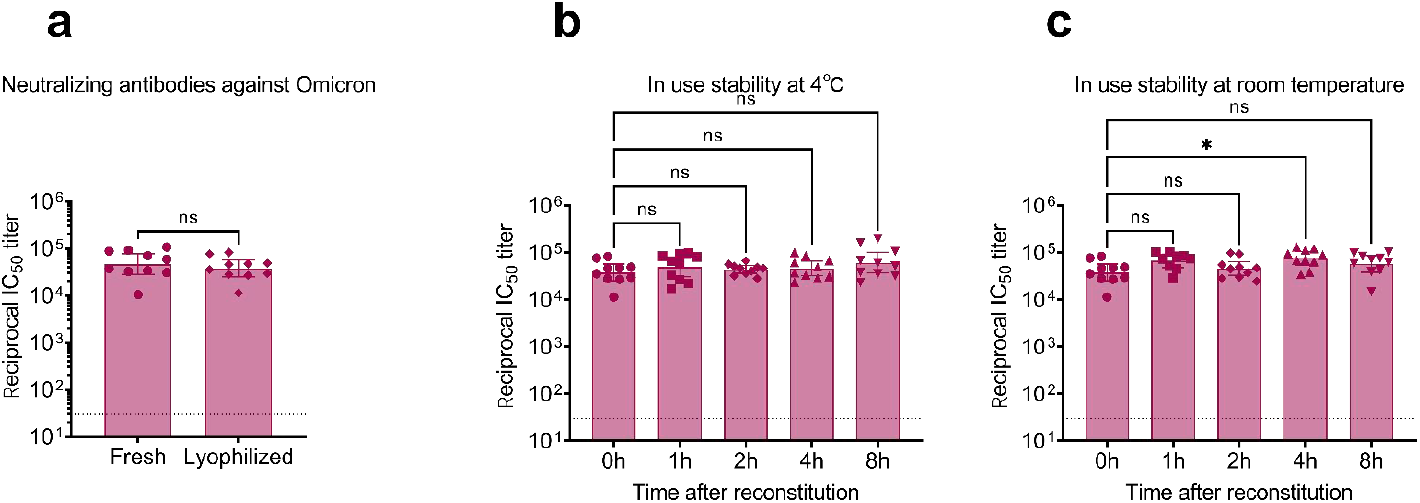
In use stability of LyomRNA-Omicron. / BALB/c mice(n=10) were immunized at a dose of 5 μg, with 2 shots administered at an interval of 21 days. Blood at 14 days post the 2^nd^ immunization was separated for detection of pseudovirus neutralizing antibodies. (a) Changes in immunogenicity after freeze drying. (b) Changes over time in immunogenicity of thawed vaccine at 4°C. (c) Changes over time in immunogenicity of thawed vaccine at ambient temperature. (a) Groups were compared by two-sided Mann-Whitney test. (b-c) Comparisons of time points were made by Kruskal-Wallis ANOVA with Dunn’s multiple comparisons test. ns=no significant, * = p < 0.05. Data are presented as geometric mean ± 95% confidence interval.

**Extended Data Fig. 10.**
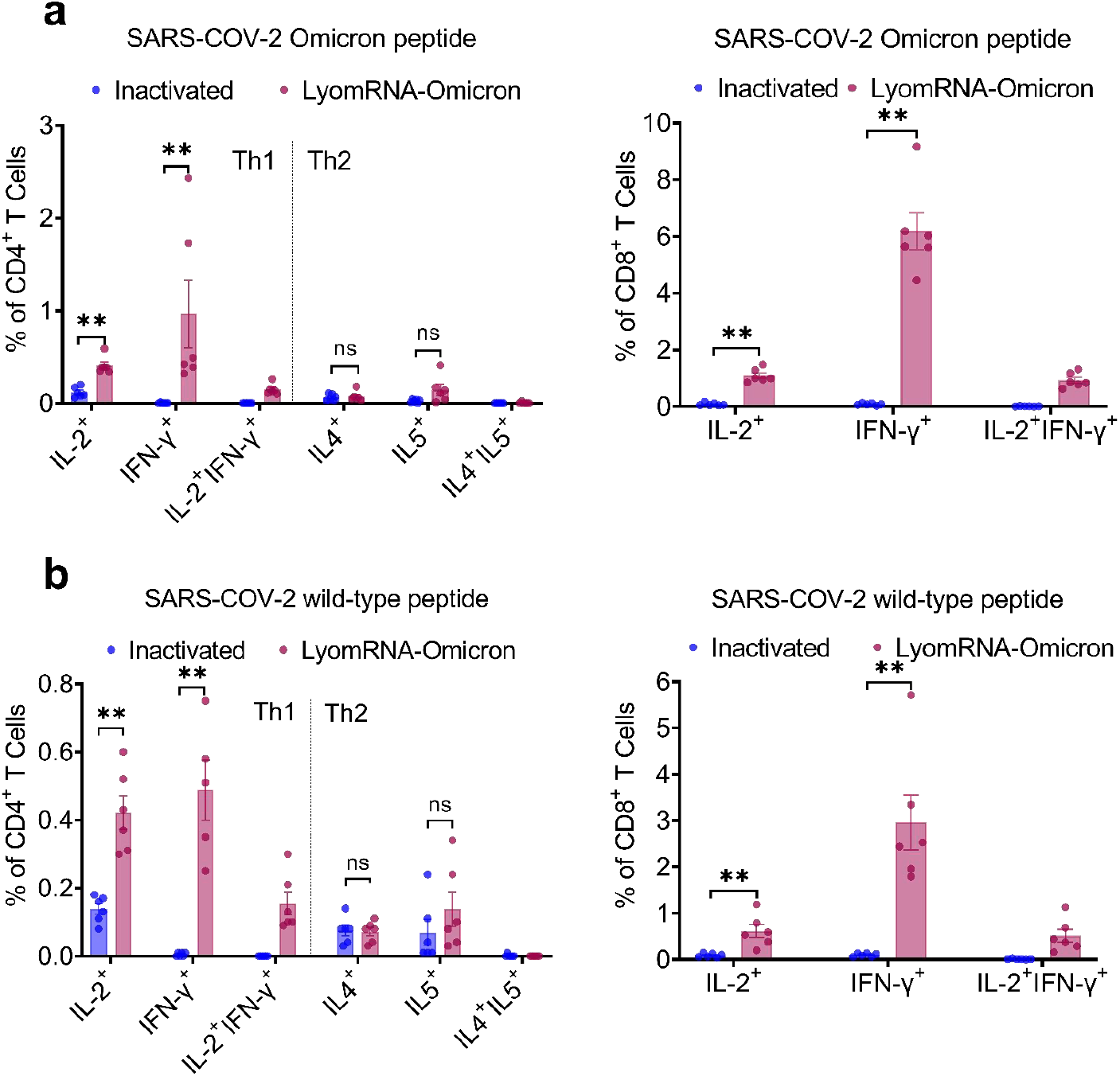
LyomRNA-Omicron-induced cellular immune response in BALB/c mice. C57BL/6N mice were vaccinated with LyomRNA-Omicron 5 μg or inactivated vaccine 0.65 U at an interval of 21 days. The spleens were obtained 28 days after the second shot for ICS. The frequencies of IL2/IFN-γ/IL4/IL5 positive CD4 T cells and IL2/IFN-γ positive CD8 killer cells specific for SARS-CoV-2 Omicron NTD-RBD peptide (a) or wild-type NTD-RBD peptide (b) were detected. Vaccine groups were compared by two-sided Mann-Whitney test. ns=no significant, ** = p< 0.01. Data are presented as mean with SEM.

**Extended Data Fig. 11.**
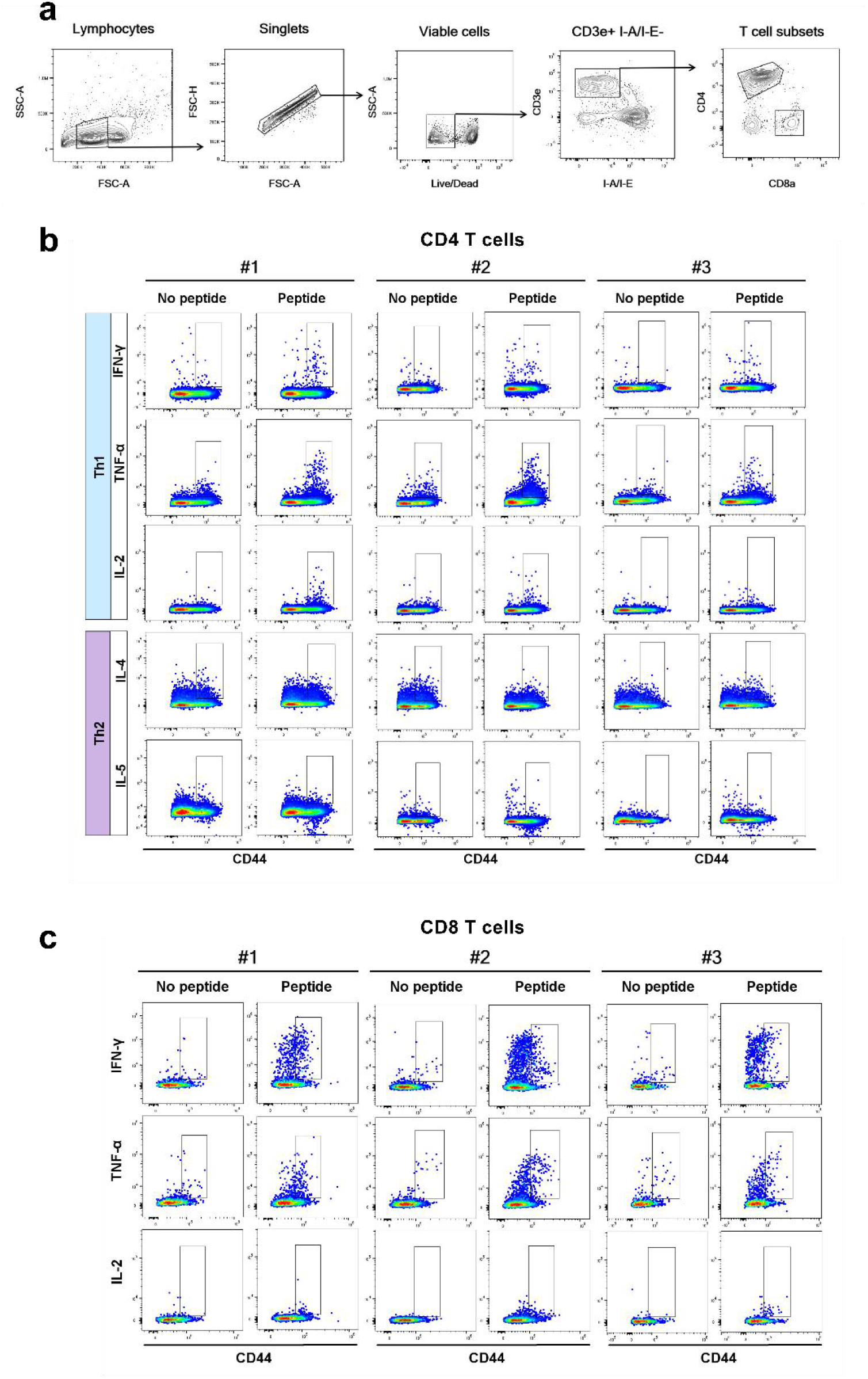
Flow cytometry panel to quantify Omicron NTD-RBD peptide specific T cells in BALB/c mice. (a) Hierarchical gating strategy to identify single, viable CD4+ and CD8+ T cells. (b-c) Gating summary of Omicron NTD-RBD peptide specific cytokine+CD44hi+ CD4+ (b) and cytokine+CD44+CD8+ (c) T cells elicited by 5 µg LyomRNA-Omicron. #1/#2/#3 represents individual mouse.

**Extended Data Fig. 12.**
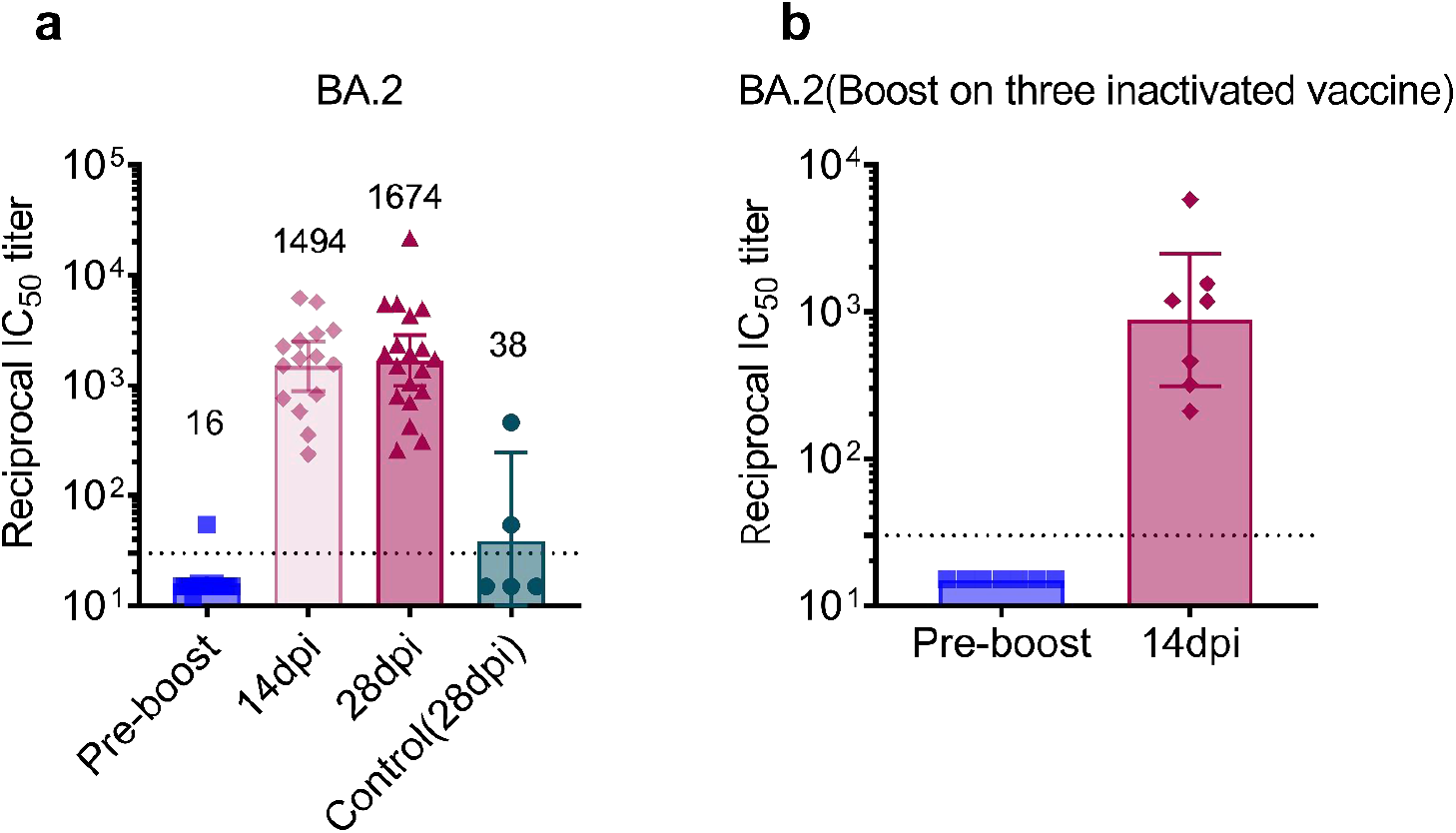
Plasma neutralizing antibodies against Omicron-BA.2 pseudoviruses following booster immunization with LyomRNA-Omicron. Omicron-BA.2 VSV-pseudoviruses from a laboratory different from those in Figure 5 were used for detection of the titers of serum neutralizing antibodies. (a) Heterogeneous booster immunization with LyomRNA-Omicron (50 μg) following two doses of inactivated vaccine. (b) Heterogeneous booster immunization with LyomRNA-Omicron (50 μg) following three doses of inactivated vaccine. Data are presented as geometric mean ± 95% confidence interval. Horizontal dashed line marks the LLOD. Values below the LLOD are set to 1/2 LLOD.

**Extended Data Fig. 13.**
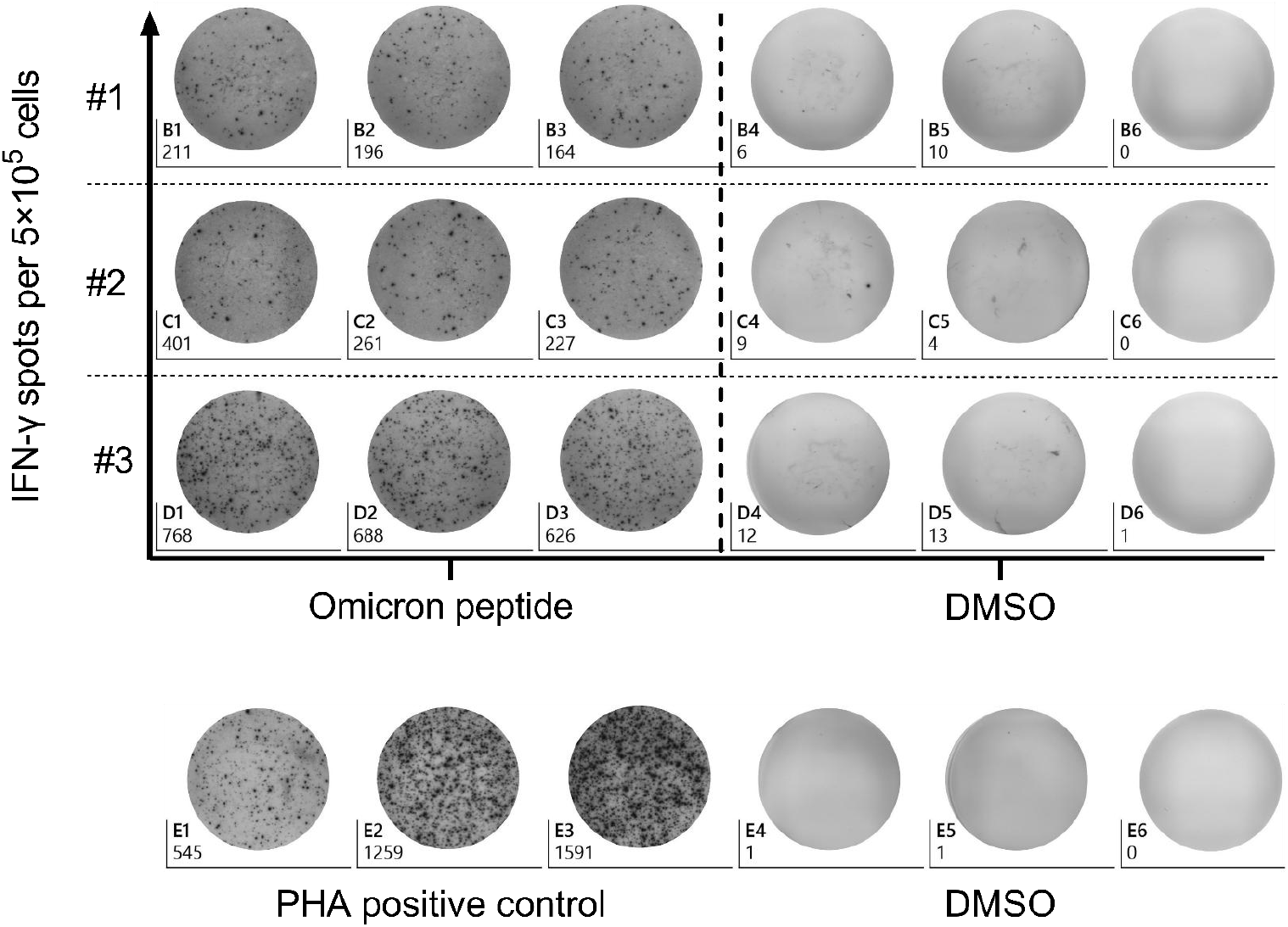
Cellular immune responses in humans following LyomRNA-Omicron booster immunization. #1/#/2/#3 represent 3 individual volunteers. Booster shot was given with 50 μg mRNA following two doses of inactivated vaccine. PBMCs (5×10^5^ cells per well) were collected #1 at day 59, #2 day 58, and #3 day 21 post booster immunization. The number of IFN-r spots was detected by ELISpot. Omicron NTD-RBD Omicron peptide or DMSO were used for stimulation in #1/#2/#3 and 5×10^5^ cells per well were used in the PHA group.

